# An environment-responsive SCLA-type GRAS protein controls reproductive strategy in *Marchantia polymorpha*

**DOI:** 10.1101/2024.11.08.622648

**Authors:** David J. Hoey, Philip Carella, Weibing Yang, Sebastian Schornack

## Abstract

By controlling their reproductive strategies, plants strike a balance between generating genetic diversity or maintaining stability, promoting long-term survival in a range of environmental conditions. The liverwort *Marchantia polymorpha* propagates clonally through vegetative gemmae or via sexually generated spores, but our understanding of the genetic pathways underpinning reproductive decision-making is limited. GRAS transcriptional regulators are highly conserved proteins across land plants, with diverse functionality in stress and development. Here, we discover a central role for the SCLA-type transcription regulator Mp*GRAS7* in both stress signalling and reproduction in Marchantia. Mp*GRAS7* is highly expressed in gemma cups and gametangiophores and is activated in response to Far Red (FR) light and abscisic acid (ABA) environmental cues. Genetic dissection further suggests Mp*GRAS7* is required to maintain gemma dormancy, gemma cup frequency, and FR-induced gametangiophore development. The sequence conservation and expression patterns of Mp*GRAS7* orthologs in flowering plants hints towards the broader function of SCLA-type GRAS proteins in stress-regulated developmental transitions in plants.

## Introduction

Plants have a remarkable ability to adjust their growth, development, and reproductive strategies to a changing local environment. In angiosperms, control of flowering time in response to environmental cues is well studied in the model plant *Arabidopsis thaliana*. In the Arabidopsis Col-0 ecotype, long days, heat stress (in long days) and drought stress accelerate flowering; while short days, cold stress, and salt stress can delay flowering (Kazan & Lyons, 2015). Some plants, such as the liverwort *Marchantia polymorpha*, have developed additional asexual (vegetative) reproduction strategies to increase the dissemination of genetic material (Ishizaki *et al*., 2016). *M. polymorpha* reproduces vegetatively via clonal gemmae produced in gemma cups, and sexually via gametes produced in gametangia which develop on umbrella-shaped gametangiophores. *M. polymorpha* is dioecious, with male organs being known as antheridiophores, and female organs as archegoniophores (Ishizaki et al., 2016), lending itself to the study of both vegetative and generative developmental processes. Recent ecological data has shown that *Marchantia polymorpha* reproductive structures show a degree of seasonality, propagating mainly vegetatively through the winter, with sexual reproductive organs emerging in the spring and summer. Male organs tend to emerge first, and female organs tend to emerge later, persisting longer as the sporophyte stage develops (Duckett *et al*., 2024). Little is known about molecular components that balance between sexual and asexual reproductive strategies.

Plants modulate their growth and development via transcriptional regulators such as the GRAS domain family named after the first well characterised proteins (GAI, RGA, SCR). GRAS transcription factors are a highly conserved family of plant proteins with diverse roles across land plants. In angiosperms, genetic studies have demonstrated key roles for GRAS domain proteins in processing environmental stimuli, for example in hormone signal transduction (e.g., gibberellic acid, GA, Silverstone *et al*., 1997). They also have diverse roles in stress and development, as well as in plant-microbe interactions (Jaiswal *et al*., 2022). Among the 19 sub-families of GRAS-domain proteins found in land plants, our understanding of the Scarecrow-Like A (SCLA) sub-family remains minimal. With a reduced set of transcriptional regulators, the liverwort *M. polymorpha* has been of particular interest to plant biologists as a tractable model system to investigate the role of transcriptional regulators in growth and development (Bowman *et al*., 2022). MpGRAS7 is the sole member of the SCLA group in *M. polymorpha,* making it an attractive target for functional genetics studies of this sub-family.

Genetic approaches have unravelled some of the regulators controlling reproduction in response to environmental cues in *M. polymorpha.* Like *Arabidopsis thaliana*, *M. polymorpha* requires long day conditions for the initiation of sexual reproduction. In short days, the plants produce more gemma cups (Benson-Evans, 1961; Kubota *et al*., 2014). The protein complex formed by Mp*GIGANTEA* (Mp*GI*) and Mp*FLAVIN-BINDING KELCH REPEAT F-BOX1* (Mp*FKF1*) are responsible for the integration of day length as a cue for gametangiophore initiation and are both essential for gametangiophore development (Kubota *et al*., 2014). Initiation of sexual reproduction in *M. polymorpha* requires not only long days but a period of continuous irradiation with Far-Red (FR) light. The FR-light responsiveness of gametangiophore initiation is dependent on PHYTOCHROME INTERACTING FACTOR (Mp*PIF*) and is a part of the Far-Red High Irradiation Response (FR-HIR) (Inoue *et al*., 2019). In *M. polymorpha*, both the photoreceptor phytochrome (Mp*PHY*) and Mp*PIF* exist as single-copy genes (Bowman *et al*., 2017) and MpPIF activates the FR-HIR response, whereas angiosperm PIFs generally function as repressors (Inoue *et* al., 2019).

Several mutants display defects in the different stages of gametangiophore development. These include gametangia differentiation and gametangiophore initiation (Yamaoka *et al*., 2018, Gutsche *et al*., 2023), and the folding of the gametangiophore stalk (Briginshaw *et al*., 2022). GA biosynthesis mutants are also impacted in gametangiophore initiation (Sun *et al*., 2023). The GRAS transcription factor Mp*DELLA* negatively regulates gametangiophore development in *Marchantia polymorpha*, as overexpression lines cannot produce gametangiophores (Hernández-García *et al*., 2021). The interaction of MpDELLA with MpPIF (Hernández-García *et al*. 2021; Briones-Moreno *et al*., 2023) provides a potential explanation for this phenotype.

Key regulators in gemma cup development and dormancy have also recently been identified. Mp*GEMMA CUP ASSOCIATED MYB-1* (Mp*GCAM1*) is critical for the formation of gemma cups (Yasui *et al*., 2019; Aki *et al*., 2022), and Mp*KARAPPO* is essential for the development of gemmae within the cup (Hiwatashi *et al*., 2019). Karrakin signalling negatively regulates the number of gemmae in a cup (Mizuno *et al*., 2021; Komatsu *et al*., 2023). Several hormone signalling pathways impact gemma cup dormancy in Marchantia. A key regulator of this is the drought-response hormone abscisic acid (ABA), analogous to seed dormancy in seed plants. Mutants in Mp*ABI3a*, a major regulator of ABA responses, germinate prematurely in the cup (Eklund *et al*., 2018). A single-copy cryptochrome in *M. polymorpha*, Mp*CRY,* is implicated in regulating gemma cup dormancy, also via ABA signalling (Liao *et al*., 2023). In addition, Auxin and ethylene are also associated with gemma dormancy (Eklund *et al*., 2015; Kato *et al*., 2015; Li et al., 2020; Kato *et al*., 2022).

Here, we identify MpGRAS7 as a key regulator of asexual and sexual reproductive structure development in liverworts. Through promoter-reporter fusions and mutant studies, we demonstrate that MpGRAS7 is necessary for appropriate gemma cup initiation, gemma dormancy, and FR-induced gametangiophore development. We further show that MpGRAS7 is a component of FR-light and ABA signalling pathways in *M. polymorpha*, where its expression is coordinated via its upstream promoter. We propose that MpGRAS7 contributes to a stress-induced shift from asexual reproduction to sexual reproduction in Marchantia coordinated via FR-light and ABA hormone cues. This study provides new insights into the function of SCLA-type GRAS transcriptional regulators. Future studies will address conserved roles and additional functions of SCLA-type GRAS proteins across land plants.

## Results

### SCLA genes are widely conserved across land plants with lineage-specific losses

First, we surveyed SCLA prevalence across all major land plant lineages and determined that they often exist as single-copy or two-copy genes (Fig. 1A). Notable exceptions to this pattern were large proliferations in certain fern lineages (*Actinostachys digitata, Alsophila spinulosa, Ophioglossum reticulatum*) and absence from others (*Azolla filiculoides*, *Salvinia cucullata*, *Ceratopteris richardii*). SCLAs were absent from all Brassicaceae surveyed, but were present in the related species *Carica papaya*, a member of the Brassicales-Malvids. Likewise, SCLA is lost from all Pinaceae and Gnetaceae species surveyed, but present in the related gymnosperm *Ginkgo biloba* (Fig. 1B). Additional losses in angiosperms were observed in *Cucumis sativus* (Cucurbitaceae) and *Liriodendron chinense* (Magnoliaceae), indicating that SCLA has been repeatedly lost across land plants.

**Figure 1.**
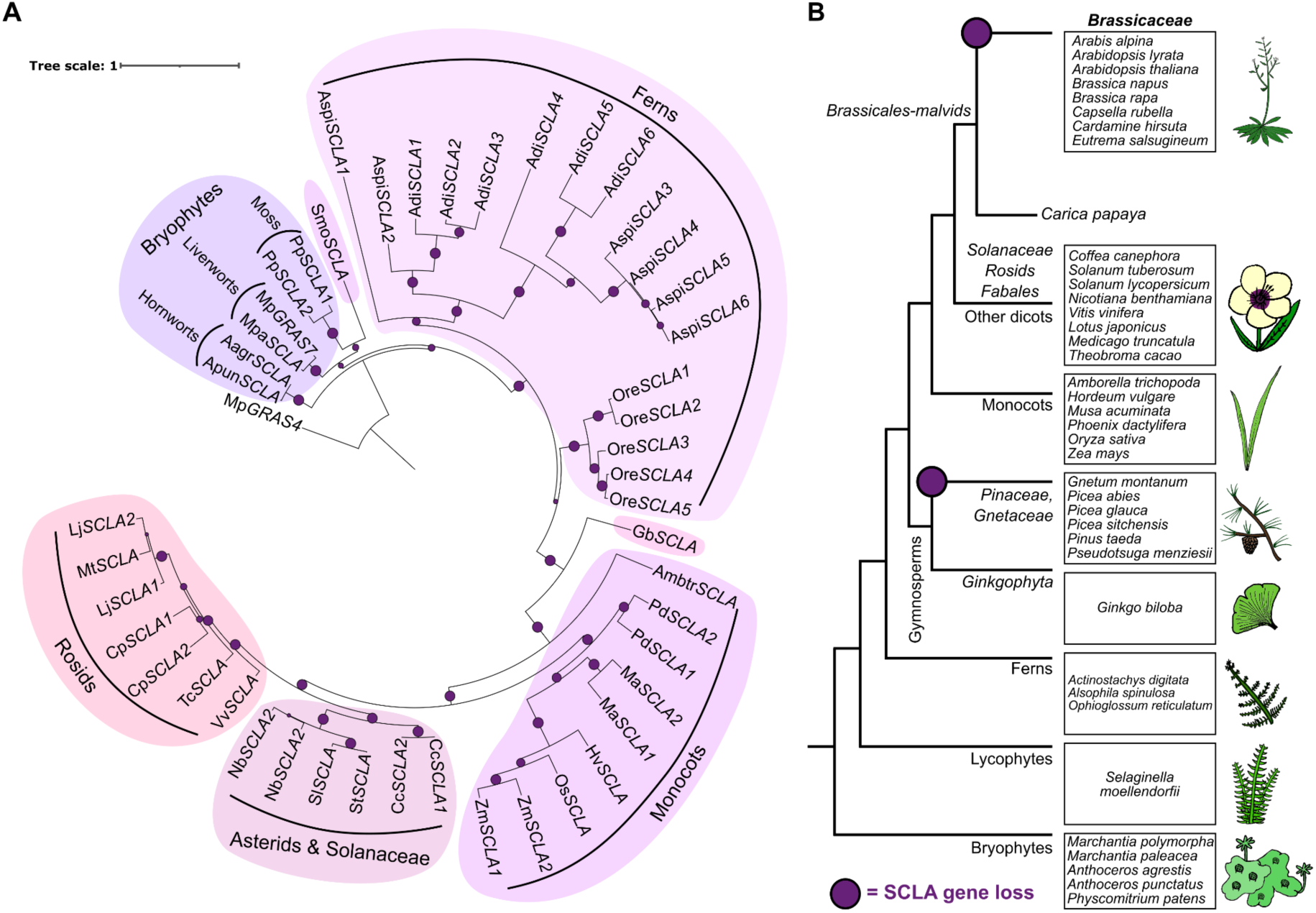
The SCLA family is conserved across land plants, although lost in several lineages including *Brassicaceae.* **A)** Maximum-likelihood phylogenetic tree of the SCLA sub-family of GRAS domain proteins. Mp*GRAS4/*Mp*LAS* used as an outgroup. Relative size of purple circles represents bootstrap support values. **B)** Graphical tree of land plant lineages which have lost the SCLA sub-family of GRAS transcription factors. SCLAs have been lost in in sequenced Pinaceae and Gnetaceae genomes (*Gnetum montanum, Picea abies, Picea glauca, Picea sitchensis, Pinus taeda, Pseudotsuga menziesii*) and *Brassicaceae* (*Arabis alpina, Arabidopsis lyrata, Arabidopsis thaliana, Brassica napus, Brassica rapa, Capsella rubella, Cardamine hirsuta*). SCLAs are present in species neighbouring these lineages, such as *Ginkgo biloba*, and *Carica papaya* respectively. Purple circles represent postulated historical losses of SCLA.

### Mp*GRAS7* is expressed in reproductive organs and responds to abiotic cues

To interrogate the role of SCLA in vegetative and sexual reproduction, we transformed *Marchantia polymorpha Tak-1* and *Tak-2* with a transcriptional reporter consisting of a 2.8 kb sequence upstream of the Mp*GRAS7* gene start codon, fused to either a glucuronidase A (GUS) reporter (*pro*Mp*GRAS7::GUS*), or a fluorescent nuclei reporter (*pro*Mp*GRAS7::tdTomato-NLS*) gene. Plants carrying the promoter-reporter constructs showed prominent reporter activity in gemma cups and gametangiophores, organs linked to vegetative and sexual reproduction, respectively (Fig. 2, Fig. S1). This included gemmae (Fig. S1A), early developing cups, mature dormant gemma cups, but not germinated gemma cups (Fig. 2A-B), as well as antheridia (Fig. 2C, Fig. S1B-D) and archegonia (Fig. 2D, Fig. S1D). *In-situ* hybridisation experiments showed a sectorised expression of Mp*GRAS7* within antheridia (Fig. 2E, Fig. S1E-G). RT-qPCR confirmed transcript enrichment in gametangiophores, with a higher expression in antheridiophores (Fig. S1H).

**Figure 2.**
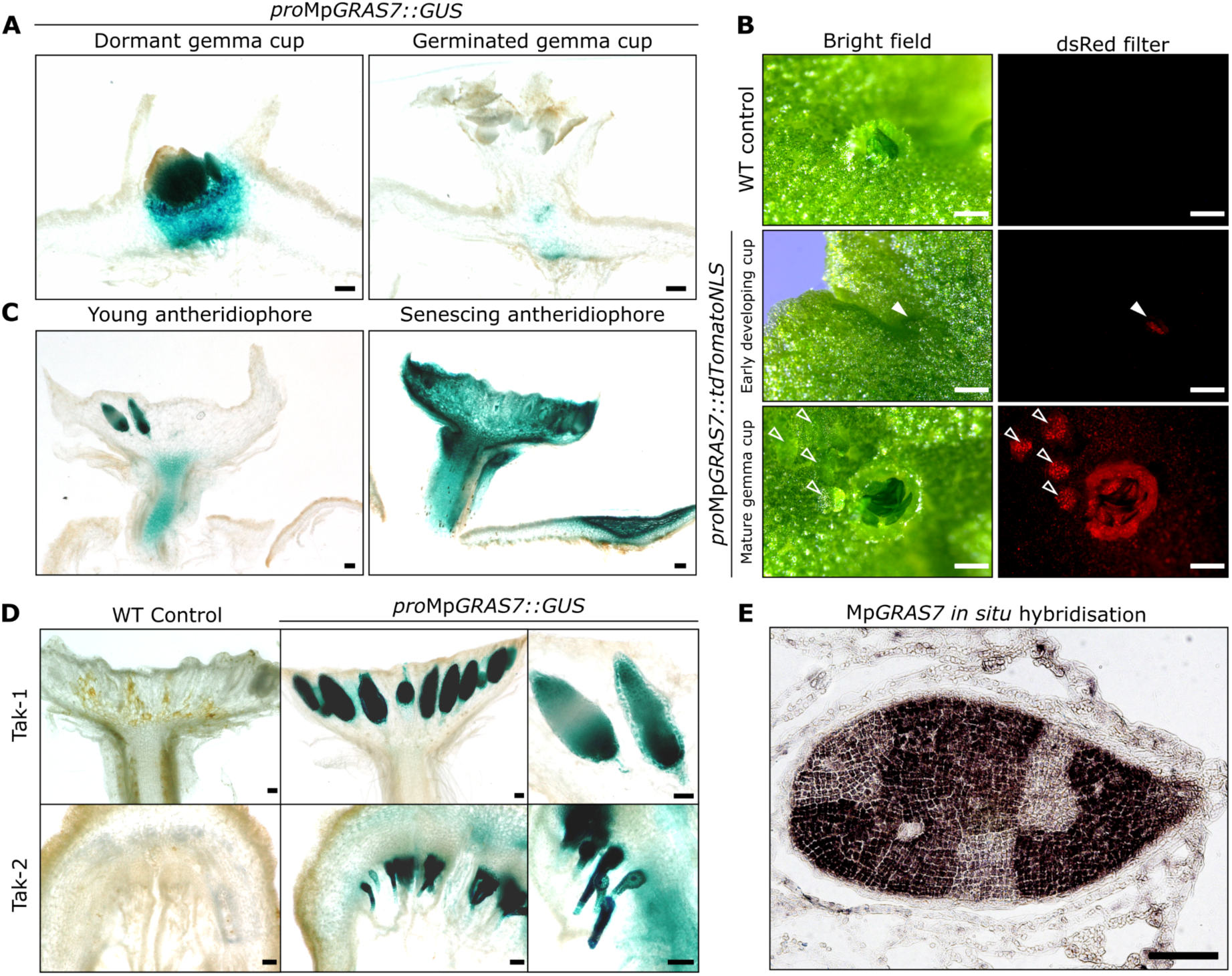
Mp*GRAS7* is expressed in reproductive organs. **A)** Agarose sections of GUS stained promoter-reporter lines, *pro*Mp*GRAS7::GUS*, showed expression in mature gemma cups but not cups which had germinating gemmae. Three independent transgenic lines were tested and all showed GUS staining in the cup. GUS staining was repeated twice with similar results. Scale bars = 200 μm. **B)** Epifluorescence imaging of *pro*Mp*GRAS7::tdTomato-NLS*, showed high expression in early developing gemma cups (filled arrow) and dormant gemmae (open arrows). Epifluorescence of gemma cups was observed in three independent experiments after three weeks of growth from gemmae. Scale bar = 1000 μm. **C)** Agarose sections of *pro*Mp*GRAS7::GUS* antheridiophores. Young antheridiophore shown is six weeks post induction, senescing antheridiophore is ten weeks post induction with FR light. Three independent transgenic lines were tested and all showed GUS staining in the antheridiophore. GUS staining of antheridiophores was repeated three times with similar results. Scale bars = 200 μm. **D)** Agarose sectioning of GUS stained *pro*Mp*GRAS7::GUS* plants in both the male (*Tak-1*) and female (*Tak-2*) backgrounds. Scale bars = 100 μm. **E)** *in situ* hybridisation of Mp*GRAS7* transcripts in an antheridium. Scale bar = 100 μm.

Since control of vegetative and sexual reproduction organ development has been previously linked to abiotic environmental cues such as FR light and ABA (Eklund *et al*., 2018; Inoue *et al*., 2019), we quantified Mp*GRAS7* transcripts via RT-qPCR and assessed the Mp*GRAS7* promoter activity in thalli of *pro*Mp*GRAS7::GUS* reporter lines upon exposure to these reproduction-relevant abiotic cues. Mp*GRAS7* upregulation was observed upon extended Far-Red light irradiation, but was absent in *gras7-1^Tak-1^* and *pif^ko^* mutants (Fig. 3A, Fig. S2A-C). Mp*GRAS7* is also activated upon humidity-stress treatment (Fig. 3C) and application of abscisic acid (ABA) (Fig. 3B,D; Fig. S3A-C). We found this activation to be dependent on the ABA receptor Mp*PYL1* (Fig. S3C). Collectively, our data suggest that Mp*GRAS7* is a component of the Far-Red High Irradiance Response (FR-HIR) and of ABA-dependent stress signalling.

**Figure 3.**
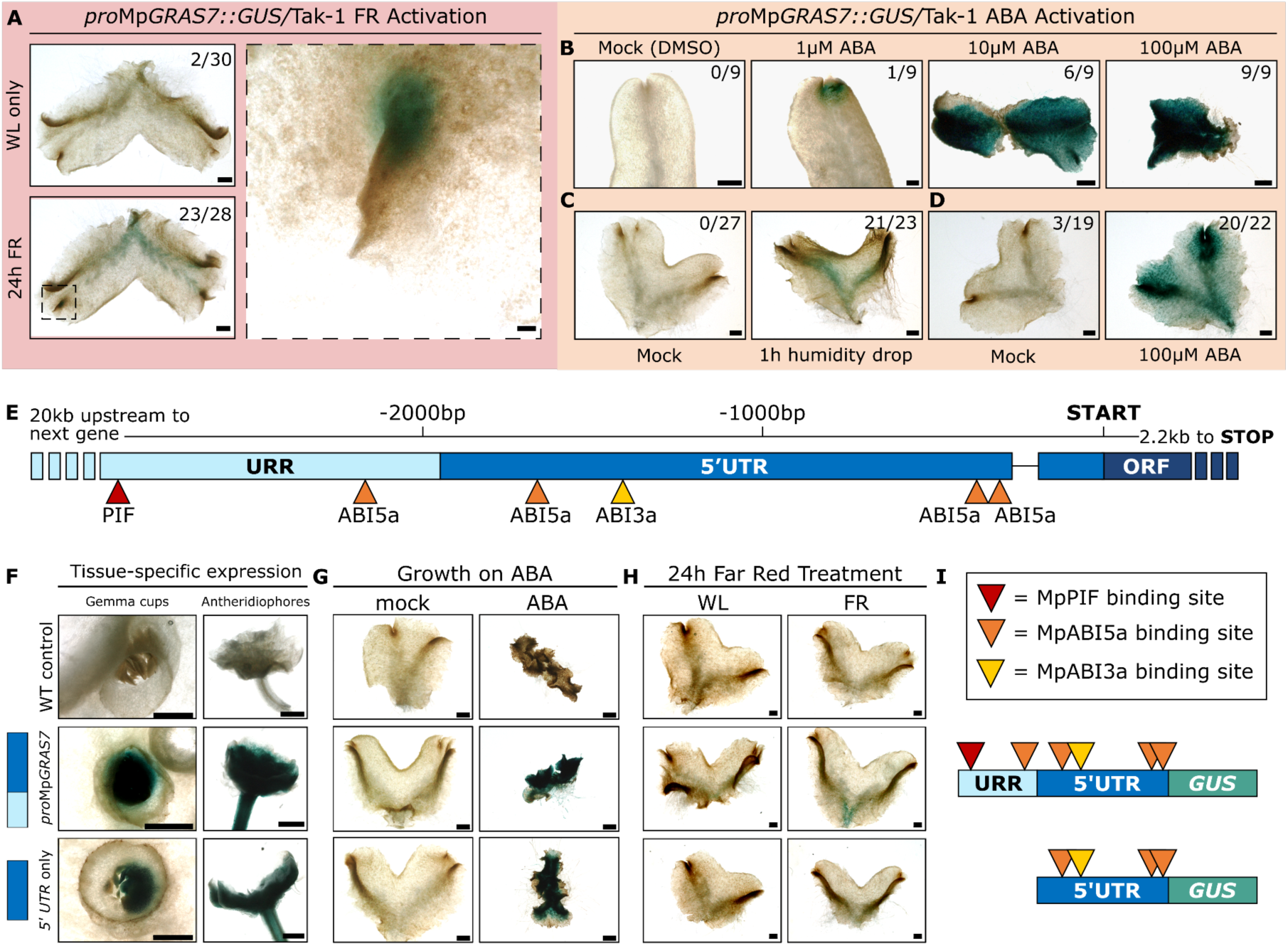
Mp*GRAS7* is activated by FR and ABA environmental cues, which target the promoter. **A)** Light microscopy images of 2-week-old GUS-stained *pro*Mp*GRAS7::GUS* plants at the 24 h FR treatment time point. WL = white light control, plants untreated with FR light. Scale bars = 1000 µm. *n=16* individual plants per condition. Inset shows activation of *pro*Mp*GRAS7::GUS* in the meristem after 24 h FR light treatment. Scale bar for inset = 100 µm. **B)** *pro*Mp*GRAS7::GUS* lines subjected to either a mock (low concentration of DMSO, equivalent to the 100 µM ABA treatment) treatment or ABA treatment via growth media. Plants were harvested after 4 weeks’ growth. 3 independent transgenic lines were tested, using 3 individual plants per line per treatment. **C)** 2-week-old *pro*Mp*GRAS7::GUS* plants treated with a humidity drop for 1 h, and GUS stained 4 hpt. *n=16* individual plants per condition. ‘Lobes with blue GUS’ were counted after GUS staining, as plants normally separate in tissue processing (numbers shown in image corners in A-D). Scale bars for B-D = 1000 µm. **D)** 2-week-old *pro*Mp*GRAS7::GUS* plants subjected to droplet-treatment with either a mock solution (low concentration of DMSO, diluted equivalent to the 100 µM ABA treatment) or ABA solution, and harvest 4hpt. **E)** Graphical representation of the upstream regulatory sequence of Mp*GRAS7* (*pro*Mp*GRAS7*). Light blue depicts a 1-kb non-transcribed Mp*GRAS7* promoter region (termed upstream regulatory region, URR). Dark blue depicts the 1.8 kb 5’-untranslated region (UTR). An intron in the 5’UTR is represented with a horizontal black line. The open reading frame (ORF) is shown in darker blue. Selected transcription factor binding sites as predicted by PlantRegMap (Tian *et al.,* 2020) (*p*<0.001) are depicted by coloured triangles (Mp*PIF* = red, Mp*ABI5a* & Mp*ABI3a* = orange). **F-H)** Promoter truncation experiments. The first construct shown is the full-length promoter (*pro*Mp*GRAS7::GUS*, followed by a construct which contains GUS driven by the 5’UTR-only (Mp*GRAS7-5’UTR::GUS*). **F)** GUS stained gemma cups (4-weeks post plating, *n*=10-15 cups per genotype) and late-stage whole antheridiophores (10 wpi, *n*=40-60 antheridiophores per genotype). **G)** GUS-stained 2-week-old plants grown on either mock media or media containing 100 μM ABA, *n*=6-9. **H)** GUS-stained 2-week-old plants after treatment with either WL or 24 h FR light, *n*=14-17. Experiments on truncated promoter lines were repeated on 2-3 independent transgenic lines. Scale bars for F-H = 1000 µm. **I)** Graphical representation of constructs used in promoter dissection experiments, colour scheme as in E.

Consistent with its Mp*PIF-* and Mp*PYL1*-dependent induction, the Mp*GRAS7* upstream regulatory region (URR) contains a predicted PIF binding motif alongside binding sites for ABI5, a transcription factor of the ABA signalling pathway. In addition, three ABI5 and one ABI3 binding sites were predicted in the Mp*GRAS7* 5’-untranslated region (5’-UTR) (Fig. 3E). GUS reporters driven by the 5’-UTR only (Mp*GRAS7-5’UTR::GUS*) were sufficient for activation of GUS signal in antheridiophores, gemma cups, and during ABA treatment (Fig. 3F-G) but did not show FR-induced GUS activity (Fig. 3H). The URR harbours the putative MpPIF binding site (Fig. 3I), which may explain the absence of a GUS signal in constructs lacking this region. We conclude that MpGRAS7 responsiveness to different abiotic cues are mediated by distinct regions of the MpGRAS7 promoter.

Looking beyond Marchantia, we find that the upstream regulatory regions of SCLA genes from bryophytes to angiosperms harbour putative transcription factor binding sites related to FR light and ABA signalling (Fig. S4A-C). Notably, *pro*Mp*GRAS7* has an exceptional proliferation of ABI3/5 binding sites, possibly reflecting its water-dependent lifestyle. Tissue-specific expression data indicate that SCLAs are particularly enriched in seeds, anthers, and silks in monocots (Fig. S4D-E), and the rice ortholog, *OsSCLA*, is responsive to ABA treatment (Fig. S4F). A comprehensive understanding of SCLA roles across land plants requires further investigation.

### MpGRAS7 is required for an appropriate response to FR light and ABA-related stress

To further understand the role of Mp*GRAS7* we generated three independent mutant alleles through CRISPR/Cas gene editing (*gras7-1^Tak-1^*, *gras7-2^Tak-2^*, *gras7-3^Tak-1^*) (Fig. S5A-D). Male *gras7-1^Tak-1^* mutants retained normal thallus and antheridiophore anatomy (Fig. S6A-F). The jacket cells of *gras7-1^Tak-1^* antheridia sporadically display an aberrant cell pattern rather than a single cell file observed in wild type (Fig. S6G-H). However, these plants produced viable sperm cells as wild type female plants produce sporophytes upon treatment with mutant *gras7^Tak-1^* sperm (Cam-1 x Cam-2: 15x archegoniophores fertilised, 3 produced sporophytes. Tak-1 x Cam-2: 21x archegoniophores fertilised, 2 produced sporophytes. *gras7-1^Tak-1^* x Cam-2: 14x archegoniophores fertilised, 1 produced sporophytes).

To delineate the extent to which downstream transcriptional responses to FR and ABA depend on Mp*GRAS7,* we compared the transcriptomes of wild type and *gras7-1^Tak-1^* mutant (2-week-old) thallus exposed to either FR light (24 h post treatment) or ABA (1 h or 4 h post treatment). Comparisons between mock and FR-treated samples revealed that a third (160/503, 31.81 %) of the wild type FR-responsive genes (Fig. 4A-C, Table S1C-D), and around a fifth (16/72, 22.22 % at 1hpt; 62/357, 17.37 % at 4hpt) of the wild type ABA-responsive genes were no longer responsive in the *gras7-1^Tak-1^*mutant (Fig. 5A-E, Tables S1D-H). A further 42 FR-responsive genes (Table S1I-K), and 14 ABA-responsive genes (Table S1L-P) were found to be significantly (p-adj<0.01) differentially expressed when directly comparing wild type and *gras7-1^Tak-1^* samples.

**Figure 4.**
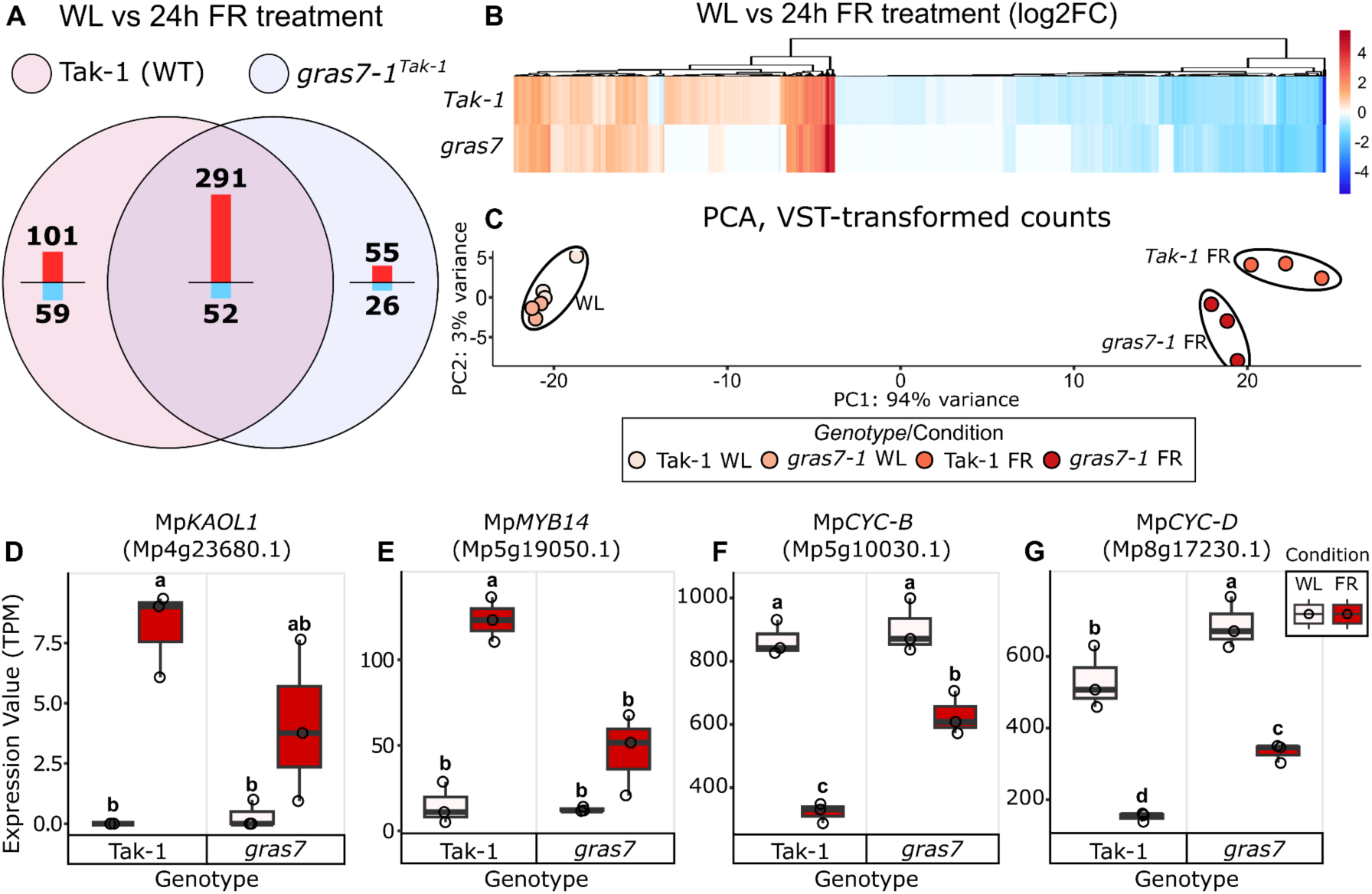
Mp*GRAS7* is required for a full transcriptional response to FR light-treatment. **A)** Venn diagram of WL vs FR-treated differentially expressed genes in different genotypes and time points. Genes were filtered by an adjusted p-value of 0.01 and Log2FC in expression of =>2 and =<2 used as cut-offs. **B)** Heat map showing all significant (p-adj<0.01) Log2FC values in response to FR light in WT and *gras7-1^Tak-1^* samples. **C)** PCA plot of variance stabilised transcripts of samples in the FR-treatment experiment. Samples mainly group by treatment, WL samples cluster closely together, whereas there is some divergence between WT and *gras7-1* FR-treated samples. **D-G)** Expression values (Transcripts Per Million, TPM) (*n*=3) of **(D)** Mp*KAOL1* (Mp4g23680.1), **(E)** Mp*MYB14* (Mp5g19050.1), **(F)** Mp*CYC-B* (Mp5g10030.1), **(G)** Mp*CYC-D* (Mp8g17230.1) from the FR-treatment transcriptome. Statistical groupings are Tukey’s HSD Post-hoc analysis (*p*<0.05).

**Figure 5.**
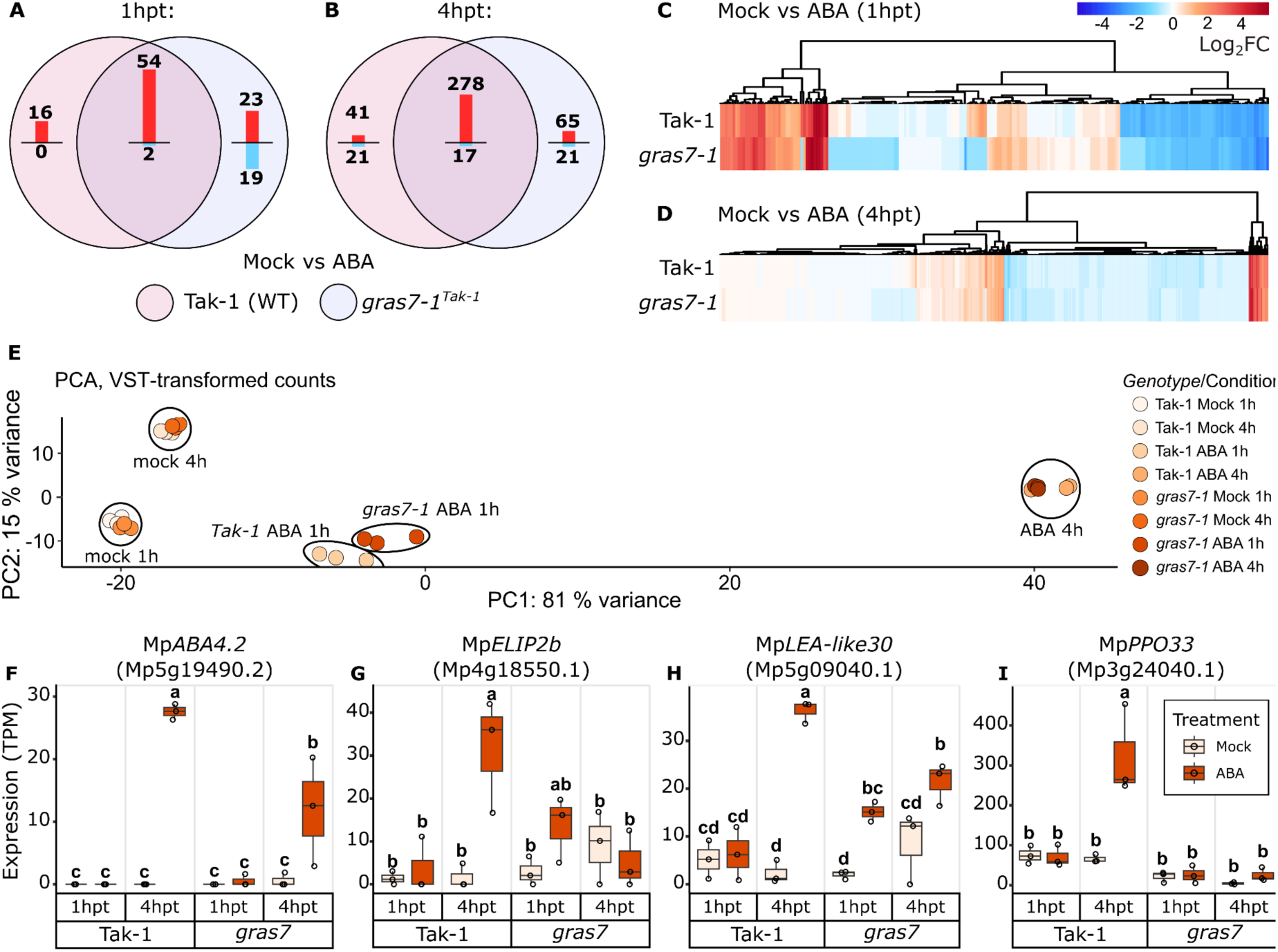
Mp*GRAS7* is required for a full transcriptional response to ABA-treatment. **A)** Venn diagram of mock vs ABA-treated differentially expressed genes in WT Tak-1 vs *gras7-1^Tak-1^* at 1h post treatment (1hpt) and **B)** 4h post treatment (4hpt). Upregulated genes are indicated in red, downregulated genes are indicated in blue. For the diagrams in A and B, genes were filtered by an adjusted p-value of 0.01 and Log2FC in expression of =>2 and =<2 as cut-offs. **C)** Heat map showing all significant (p-adj<0.01) Log2FC values in response to ABA in WT and *gras7-1^Tak-1^* samples at 1 hpt. **D)** Heat map showing all significant (p-adj<0.01) Log2FC values in response to ABA in WT and *gras7-1^Tak-1^*samples at 4 hpt. **E)** PCA plot of normalised expression of samples in the FR-treatment experiment. Samples mainly group by treatment, apart from in ABA treated samples at 1hpt. **F-I)** Expression values (Transcripts Per Million, TPM) (*n*=3) of **(F)** Mp*ABA4.2* (Mp5g19490.2), **(G)** Mp*ELIP2b* (Mp4g18550.1), **(H)** Mp*LEA-like30* (Mp5g09040.1), **(I)** Mp*PPO33* (Mp3g24040.1) from the ABA-treatment transcriptome. Statistical groupings are Tukey’s HSD Post-hoc analysis (*p*<0.05).

Among the Mp*GRAS7-*dependent FR-induced genes is the Gibberellic acid (GA)-biosynthesis gene Mp*KAOL1* (Mp4g23680.1), which catalyses GA_12_ biosynthesis in *Marchantia polymorpha* and has been shown to regulate gametangiophore development (Sun *et al*., 2023). Mp*KAOL1* is upregulated in wild type plants upon FR treatment (log_2_FC=5.36, p-adj=0.0014), but not significantly in *gras7-1^Tak-1^*mutants (Fig. 4D). Transcript levels of Mp*MYB14* (Mp5g19050.1), encoding a transcription factor involved in directing phenylpropanoid metabolic stress responses in *M. polymorpha* (Albert *et al*., 2018; Kubo *et al.,* 2018; Berland *et al*., 2019; Carella *et al*., 2019), are also significantly lower in FR-treated *gras7-1^Tak-1^* mutants compared to similarly treated wild type (Fig. 4E). The expression of two cell cycle regulators, Mp*CYC-B* (Mp5g10030.1) and Mp*CYC-D* (Mp8g17230.1) are reduced upon continuous FR exposure in wild type, but significantly less so in FR-treated *gras7-1^Tak-1^* (Fig. 4F-G). Responses to continuous FR treatment such as ABA biosynthesis and response, circadian rhythm, and other GA-biosynthesis genes were not significantly altered in the mutant.

Among the Mp*GRAS7*-dependent ABA induced transcripts is Mp*ABA4* (Mp5g19490.2), a secondary transcript encoding a neoxanthin synthase for ABA biosynthesis (Fig. 5F). Notably, the primary transcript of Mp*ABA4* remains responsive in ABA-treated *gras7-1^Tak-1^* mutants. Several ABA-response genes were no longer significantly activated in *gras7-1^Tak-1^* mutants (e.g. Late Embryogenesis Abundant Mp*LEA-like30* (Mp5g09040.1); and the Early Light Inducible Protein (ELIP) genes (Mp4g18580.1, Mp6g20760.1, Mp4g18550.1, Mp4g18590.1) (Fig. 5G-H). A specialised metabolite-regulating polyphenol oxidase gene, Mp*PPO33*, is upregulated in ABA-treated wild type samples but not activated in ABA-treated *gras7-1^Tak-1^* (Fig. 5I). Most LEA-like genes and other ABA-responsive genes such as ABA biosynthesis, dehydrins, and sucrose transport were not significantly altered in the ABA-treated mutant.

Together, the expression data suggest that Mp*gras7^ge^*plants are unable to mount an appropriate response to either FR or ABA-related stress. Consistent with a partial contribution of Mp*GRAS7* to FR and ABA responses, we found that FR- and ABA-induced growth inhibition (Fig. S7A-E) and Mp*PIF*-dependent FR-induced orthotropic growth responses (Fig. S8A-D) remain intact in *gras7-1^Tak-1^*mutants.

### Mp*GRAS7* balances vegetative and sexual reproductive organ initiation

Since the Mp*GRAS7* promoter is active in both types of reproductive organs, we investigated the development of gemma cups and gametangiophores in *gras7*^ge^ mutants. Independent CRISPR/Cas9 generated male and female mutant lines *gras7-1^Tak-1^* and *gras7-2^Tak-2^*are unaffected in their thallus growth and morphology, but develop more gemma cups compared to their wild type counterparts, which is most prominent from four to five weeks of thallus growth onwards (Fig. 6A,D). Moreover, *gras7-1^Tak-1^* and *gras7-2^Tak-2^* gemmae develop rhizoids inside the gemma cup earlier than wild type (Fig. 6B,E) suggesting that appropriate gemma dormancy is lost in *gras7-1^Tak-1^*and *gras7-2^Tak-2^* mutants. While male and female *gras7^ge^*mutants develop greater numbers of gemma cups, both mutants display a delayed onset of gametangiophore development resulting in lower overall numbers of gametangiophores at 2- and 3-weeks of growth in FR light conditions (Fig. 6C,F). Importantly, perturbations in the timing and dormancy of *gras7-1^Tak-1^* reproductive organs revert back to wild type when transformed with a construct harbouring *pro*Mp*GRAS7::GRAS7* (Fig. 6G-I). These results suggest that *gras7^ge^* mutants are shifted towards vegetative propagation. As such, Mp*GRAS7* functions as a negative regulator of gemma cup initiation, and a positive regulator of gemma cup dormancy and gametangiophore initiation.

**Figure 6.**
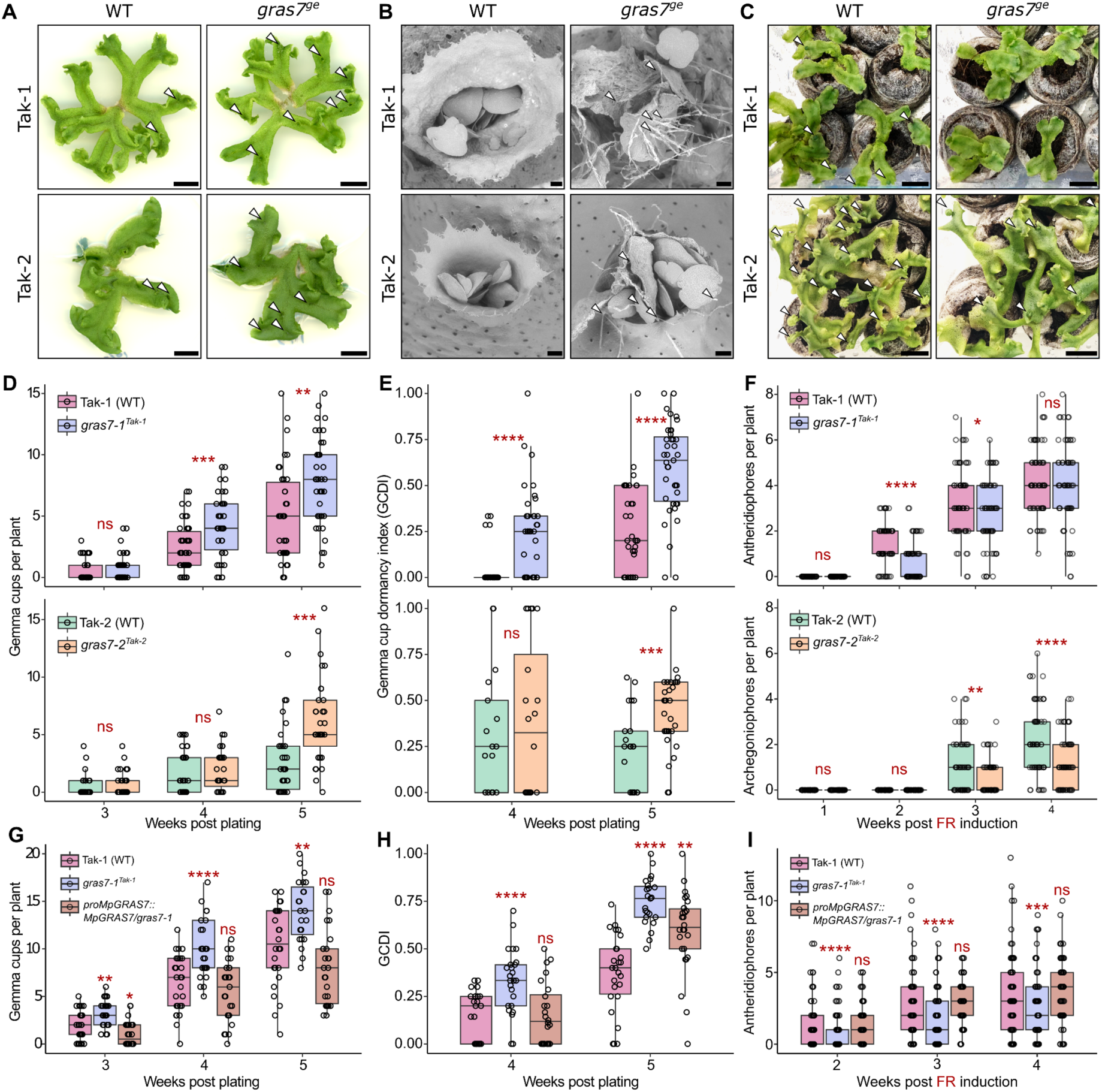
Gemma cup numbers, gemma cup dormancy, and gametangiophore initiation are shifted in *gras7^ge^* mutants. **A)** 5-week old thallus showing WT and *gras7* mutants in the male (Tak-1) and female (Tak-2) backgrounds. Mutants grow normally but develop gemma cups more frequently. White arrows indicate gemma cups. Scale = 1 cm. **B)** SEM surface imaging of WT and *gras7* gemma cups in Tak-1 and Tak-2 backgrounds at 5-weeks post-plating. White arrows denote rhizoid growth from gemmae, indicating loss of in-cup dormancy. Scale = 1000 µm. **C)** Gametangiophore emergence of WT and *gras7* mutants at 2-weeks (Tak-1) and 3-weeks (Tak-2) after induction by far-red (FR) light. Scale = 1 cm. **D)** Quantification of gemma cups per plant in WT and *gras7^ge^* mutants at 3-, 4-, and 5-weeks post-plating. Total *n*=38-42 per genotype, *N*=3 independent replicates. **E)** Gemma cup dormancy index (GCDI) of WT and *gras7* mutants at 4- and 5-weeks post plating in Tak-1 and Tak-2 mutant backgrounds, where 0=dormant and 1=not dormant. GCDI is calculated by dividing the number of gemma cups with rhizoids by the total number of gemma cups on an individual plant. Plants with zero gemma cups cannot be included in this analysis. Total *n*=38-42 per genotype, *N*=3 independent replicates. **F)** Quantification of gametangiophore emergence of WT and *gras7* mutants at 1-, 2-, 3-, and 4-weeks after induction by FR light. *gras7* mutants have later emergence of both antheridiophores (Tak-1) and archegoniophores (Tak-2). Up to 16 individuals per replicate, *N=5* replicates (total *n*=59-61 plants per genotype). **G)** Quantification of gemma cups per plant in WT, *gras7-1*, and *proMpGRAS7::GRAS7/gras7-1* at 3, 4, and 5 weeks post-plating. Total *n=*23-26 individual plants per genotype. **H)** GCDI of WT, *gras7-1*, and *proMpGRAS7::GRAS7/gras7-1* at 4- and 5-weeks post plating, calculated as in C. Total *n*=23-26 individual plants per genotype. **I)** Quantification of gametangiophore emergence of WT, *gras7-1, and proMpGRAS7::GRAS7/gras7-1* at 2-, 3-, and 4-weeks after induction by FR light. Up to 16 individuals per replicate, *N=*4-5 replicates per genotype (total *n*=62-75 plants per genotype). Statistical tests for D-I are pairwise compared to the WT control at each time point, two-tailed student’s t-test. ns = not significant, * = *p*<0.05, ** = *p*<0.01, *** = *p*<0.001, **** = *p*<0.0001.

## Discussion

This study assigns a new function to an SCLA-type GRAS protein, a GRAS sub-family with limited functional characterisation. Members of this sub-family can be identified by the presence of a specific 29 AA stretch (Cenci & Rouard, 2017). Our work demonstrates that the *M. polymorpha* transcriptional regulator Mp*GRAS7* contributes to signal transduction of the environmental cues (FR light and ABA) and controls the timing of both vegetative and sexual reproductive organ development. Given its role in balancing reproductive organ numbers and stress-responsive expression, we propose that MpGRAS7 functions as a stress-induced reproductive switch, possibly integrating environmental cues through FR light and ABA signalling. This distinguishes MpGRAS7 from other transcriptional regulators, such as MpTGA, which specifically influences gametangiophore development (Gutsche *et al*., 2023). To our knowledge, no single gene influencing both gametangiophore and gemma cup frequency in opposite ways has been identified previously.

Our Mp*GRAS7* promoter-reporter fusions defined the developmental context specific expression of Mp*GRAS7* in gemma cups, emerging gametangiophores, and gametangia within gametangiophores aligning well with public expression data (Proost & Mutwil, 2018; Yasui *et al*., 2019; Kawamura *et al*., 2022) as well as an inducible expression upon FR light or ABA treatment. The promoter used to highlight these expression patterns was also sufficient to complement the *gras7-1^Tak-1^* mutant, suggesting that it harbours the essential regulatory elements for MpGRAS7 functionality in both its responsiveness to the environment as well as its regulation of reproductive organ initiation.

MpGRAS7 likely links with other transcriptional regulators of gametangiophore development. In a yeast-two-hybrid study, Briones-Moreno *et al*. (2023) report MpGRAS7 to be a direct interactor of MpGI, MpTCP1, MpKAN, and MpDELLA, which have all been previously reported to have functions in gametangiophore development (Kubota *et al*., 2014; Busch *et al.,* 2019; Briginshaw *et al*., 2022). MpDELLA interacts with MpPIF and might function as an inhibitor of sexual reproduction based on the absence of gametangiophores in constitutive MpDELLA expression lines (Hernández-García *et al*. 2021). MpDELLA is likely to be upstream of MpPIF and MpGRAS7, but no Marchantia *della* mutants could be obtained to test this hypothesis (Hernández-García *et al*. 2021).

Current knowledge of the seasonality of the reproductive strategies of *Marchantia polymorpha* is consistent with the integration of the cues of day length (Kubota *et al*., 2014) and high FR light irradiance (Inoue *et al*., 2019). FR light is often characterised as a canopy or shade signal, but in Marchantia end-of-day or intermittent FR irradiance is insufficient for gametangiophore induction, suggesting that *M. polymorpha* initiating sexual reproduction in response to FR light is not a shade avoidance response (Inoue *et al*., 2019). Our study indicates that *M. polymorpha* may also integrate drought stress signalling into coordinating the timing of sexual initiation. This notion is reminiscent of previous findings that drought stress accelerates reproduction from studies in other model plant species (Kazan & Lyons, 2015). Given that appropriate timing of reproduction is essential for the dissemination of offspring, Mp*GRAS7* may confer a fitness benefit. This aspect could be the subject of future studies involving *gras7^ge^*mutants.

Future studies should assess whether SCLA-type GRAS transcriptional regulators fulfil conserved roles in integrating environmental cues across land plants or have evolved lineage-specific functions. Interestingly, a MpGRAS7 ortholog in *Medicago truncatula*, recently characterised to be involved in leaflet initiation (He *et al*., 2024), lacks predicted PIF and ABI3/5 binding sites. Additionally, several independent losses of SCLA-type genes across land plants suggest that SCLA regulatory activities may be either dispensable or fulfilled by other genes in some lineages.

In conclusion, our work has identified a single gene acting as a switch between the initiation of gemma cups for vegetative reproduction and gametangiophores for sexual reproduction. MpGRAS7 responsiveness to FR light and drought hormone signalling enables Marchantia liverworts to strike a reproductive decision depending on a conducive environment.

## Data availability

Materials generated by this study are available from the corresponding author on request. All data are available in the main text and supplementary materials. Raw sequencing reads for the Far-Red treatment and abscisic acid treatment transcriptomes are deposited at NCBI under BioProject number PRJNA1171426.

## Author contributions

D.J.H., P.C., and S.S. designed the project. D.J.H, P.C., and W.Y. carried out experiments. D.J.H. and S.S. analysed the data. D.J.H and S.S. wrote the manuscript. All authors read and edited the manuscript.

## Ethics declarations

## Acknowledgements

The authors would like to thank all members of the Schornack group past and present for helpful discussions and critical comments. Thanks to Alan Wanke, Ed Harris, and Chetan Pandey for technical support. Thanks to Elisa Mogollón Pérez and Matthew MacLeod for regular maintenance of liverwort lines. Thanks to Jorge Hernández-García for providing the *pif^ko^* mutant and complemented line, and Daisuke Takezawa for providing the *pyl-1^ge2b^* mutant allele. This work was funded by the Gatsby Charitable Foundation (GAT3395/GLD), Royal Society (UF160413, RGF\EA\180002), BBSRC OpenPlant initiative (BB/L014130/1), and a Natural Sciences and Engineering Research Council of Canada (NSERC) postdoctoral fellowship (to P.C.).

## Materials & Methods

### Plant growth conditions

*Marchantia polymorpha* accessions Tak-1 (male) and Tak-2 (female) were grown from gemmae on half–strength MS (Murashige and Skoog) media supplemented with Gamborg B5 vitamins (Duchefa M0231, pH 6.7) and solidified with 1.5 % (w/v) Bacto™ Agar (Fisher Scientific). This media is hereafter known as ½ MS B5, and is the primary medium which plants are grown on in this manuscript. Plants were grown under continuous white light (70 *μE m^−2^ s^−1^*) at 22 °C. For growth of plants in preparation for transformation, plants were instead grown on half–strength MS (Murashige and Skoog) media (Duchefa M0222, pH 5.7) and solidified with 0.8 % (w/v) Plant Agar (Duchefa). This media is hereafter known as ½ MS Plant. For liquid culture of *M. polymorpha*, plants were grown in 1x 0M51C. 1x 0M51C is supplemented with 1 % (w/v) sucrose for *Agrobacterium* co-culture (Kubota *et al*., 2013).

To induce gametangiophore development, 2-week-old thalli were grown from gemmae in Steri-Vent (high) containers (Melford, product code: S1686) on Jiffy-7® (35 mm) peat pellets. 16 x 35 mm peat pellets per container were moved with aseptic technique using forceps, and hydrated with 125 mL sterile ddH_2_O. These were left overnight to soak. If pellets did not absorb all water, excess was poured off. If pellets were too dry, a little extra water was added (<10 mL) before placing gemmae (1x per pellet). Containers were moved from growth under white light (WL) after 2 weeks to growth under white light supplemented with FR light (wavelength 730-740nm).

### Plasmid construction and plant transformation

A 2.8 kb promoter region upstream of the start site of Mp*GRAS7* (Mp8g01770.1) was cloned from Tak-1 genomic DNA using primers with *att*L sites for cloning into pDONR-221 by BP clonase (Invitrogen) reaction. *pro*Mp*GRAS7::tdTomato-NLS* and *pro*Mp*GRAS7::GUS* vectors were created by recombination into pMpGWB316 and pMpGWB104 *Marchantia* Gateway destination vectors from Ishizaki et al., (2015) using LR clonase II (Invitrogen) according to the manufacturer’s instructions. Transformation of existing mutants *pif^ko^* (obtained from Hernandez-Garcia et al., 2021)*, pyl-1^ge2b^* (obtained from Jahan *et al*., 2019) with *pro*Mp*GRAS7::GUS* required transformation with pMpGWB304, the same GUS vector but with chlorsulfuron resistance rather than hygromycin resistance, as these mutants had existing resistance to hygromycin from CRISPR vectors.

Promoter truncations were produced by amplification of fragments directly from the *pro*Mp*GRAS7*::221 donor vector. The 5’-UTR was amplified with *att*B sites for recombination into its own pDONR-221 vectors, and then recombined into the destination pMpGWB104 vector also by LR clonase II reactions. Constructs from external sources can be found in Table S2. Constructs produced in this study can be found in Table S3. Primers used for cloning can be found in Table S4. All donor vectors produced were validated by Sanger sequencing using M13F and/or M13R primers.

To generate CRISPR constructs, sgRNAs were designed using the CRISPR direct tool (Naito et al., 2015), and oligos were synthesised according to the instructions in Sugano *et al*. (2018). sgRNA and oligo sequences can be found in Table S5 and S6, respectively. The oligos were annealed by heating to 96 °C and leaving to cool to room temperature overnight. These were then cloned into a pre-digested (PstI and SacI) entry vector pMpGE_En01 by Infusion reaction. The resulting construct was then recombined into the CRISPR destination vector pMpGE010 using LR Clonase II (Invitrogen). These constructs were all transformed into *Agrobacterium tumefaciens GV3101 (pMP90)* by electroporation, and subsequently used in transformations.

Complementation constructs were generated by establishing a Gateway destination vector with a ccdb/CmR cassette upstream of a synthetic Mp*GRAS7* sequence (herein synGRAS7), with PAM sites of sgRNA sequences removed (due continuing presence of Cas9 in the mutants). synGRAS7 was synthesized by a commercial provider (Genewiz) in a pUC backbone, with a HindIII and EcoRV immediately preceding the gene, and a SacI site immediately thereafter. A ccdb/CmR cassette was amplified by PCR from an existing Gateway vector and inserted into the EcoRV site by blunt-end ligation. The resulting plasmid was digested with HindIII and SacI, and ligated into a Marchantia destination vector (pMpGWB301), which had also been digested with HindIII and SacI. *pro*Mp*GRAS7* was added to the vector by LR recombination, resulting in a Marchantia expression vector with the native promoter driving synGRAS7. This construct was transformed into *GV3101 (pMP90)* and transformed into *gras7-1^Tak-1^*mutants.

*M. polymorpha* plants (accessions Tak-1 and Tak-2) plants were transformed by co-culturing *Agrobacterium tumefaciens* with regenerating thalli as described in Kubota *et al*. (2013). Transgenic liverworts were all selected on solidified ½ MS Plant (pH 5.6) supplemented with cefotaxime (125 μg/mL), and either hygromycin B (15-25 μg/mL) or chlorsulfuron (0.5-1 μM) in accordance with the resistance conferred by the vector. Wild type and transgenic lines used or produced in this study can be found in Table S7.

### DNA extraction and genotyping

Genomic DNA was extracted from flash frozen tissue of CRISPR candidates using a CTAB extraction method. Tissue was homogenised, then 2 mL CTAB buffer (4 % w/v CTAB, 1.4 M NaCl, 100 mM TrisHCl pH 8, 3 % w/v polyvinylpyrrolidone, 20 mM EDTA pH 8, 1 % v/v β-mercaptoethanol) was added to ∼100 mg frozen tissue per sample, and thoroughly vortexed. The homogenate was then incubated at 60 °C for 30 min, then centrifuged for 5 min at 14,680 rpm. The supernatant was then treated with RNase A, then incubated for a further 20 min at 32 °C. An equal volume of phenol-chloroform-isoamyl alcohol (25:24:1) was added to each tube and the tube vortexed. Samples were centrifuged for 1 min at 14680 rpm. The upper aqueous layer was then removed and combined with an equal volume of chloroform-isoamyl alcohol (24:1). The aqueous layer was removed and placed into a new 2 mL tube. To each sample, 0.7 volume of cold isopropanol and 0.1 volume 3 M sodium acetate (pH 5.2) was added. The samples were incubated at -20 °C overnight to allow precipitation. Samples were then centrifuged at 14,680rpm for 10 min, isopropanol was removed, and the pellet was washed with cold 70 % (v/v) ethanol twice. Pellets were dried at room temperature for 30 min or until the pellet became clear, and then suspended in 40 μL TE buffer (10 mM Tris, pH 8, 1 mM EDTA).

For genotyping, 500 ng of gDNA was digested overnight (12 hours) at 37 °C with the appropriate restriction enzyme (SacI for sgRNA1, SfcI for sgRNA2) and Cutsmart® buffer. Undigested controls were included in this incubation (gDNA with water and buffer only). PCR fragments were then amplified from undigested and digested samples, and run on a 0.8% (w/v) agarose gel. If a band was amplified from both the digested and undigested samples, it indicated a successful edit in the genome. A subset of the successful candidates was chosen, and a DNA fragment was amplified from gDNA. This fragment was then ligated into a pBlueScript backbone (pre-digested with EcoRV), validated, and three clones were sent to sequence per candidate. In all cases, mutants were considered stable when the genomic sequence was consistent between two gemma-generations. Alleles with predicted frameshift mutations resulting in a stop codon were used for further experimentation.

Genotyping of transgenic lines was completed using a Phire Plant Direct PCR Master Mix Kit (Thermo-Fisher Scientific, #F-160S). For crude extraction of gDNA, A piece of thallus (approximately 1 mm x 1 mm) was placed into a 1.5 mL Eppendorf with 100 μL of the dilution buffer provided in the kit, and ground with a pestle. The mixture was heated to 95 °C for 5 min and centrifuged for 2 min at room temperature at 12,500rpm. The supernatant was then used for PCR analysis using the Phire Direct master mix and genotyping primers.

Primers used for genotyping are shown in Table S8. Details of PCR cycles are shown in Table S9.

### Confocal, epifluorescence, light, SEM microscopy

Confocal fluorescence microscopy (Leica TCS SP8 equipped with HyD detectors) was performed as described in Carella *et al*. (2018) using tdTomato excitation = 554 nm; WL laser power = 10 %. Chlorophyll autofluorescence was imaged at the spectral range of 700-750 nm to avoid tdTomato signal. For epifluorescence images, a and epifluorescence microscopy a Leica M165FC dissecting scope was used with a dsRed filter applied for imaging tdTomato signal. Light microscopy was performed with a Keyence VHX-5000 digital microscope or an Olympus stereoscope fitted with an SLR camera. Images were processed and scale bars added with ImageJ or Inkscape. SEM surface imaging was carried out with a Hitachi Desktop SEM (TM4000Plus) with cool-stage. The cooling stage was used to prolong imaging time of each sample. When the sample was mounted, temperature was 4 °C. When the vacuum was applied, this temperature was then lowered to -20 °C, which would allow for around 10 min of surface imaging per sample.

### RNA extraction and RT-qPCR

RNA was extracted from flash-frozen liverworts using Invitrogen PureLink™ Plant RNA Reagent (Catalog number: 12322012) according to the manufacturer’s instructions for small-scale RNA extraction. DNA contamination was removed with Invitrogen TURBO™ DNase treatment (Catalog number: AM1907) treatment, in accordance with the manufacturer’s instructions. cDNA was synthesised using Invitrogen Superscript™ II Reverse Transcriptase (Catalog number: 18064071) from 2000 ng RNA. cDNA was diluted tenfold for qPCR analysis. Expression of genes of interest in *M. polymorpha* is normalised to housekeeping genes Mp*EF1a* and Mp*ACT* (Table 19, Saint-Marcoux *et al*., 2015). All RT-qPCR experiments were subject to the following temperature cycling conditions: 95 °C 5 min, [95 °C 20 s, 50 °C 20 s, 72 °C 20 s] x45, followed by melt curve analysis 50 °C to 97 °C. Melt curve analysis was used to ensure only one PCR product was being amplified. All RT-qPCR experiments were carried out alongside a ‘no RT’ control, a sample of RNA from the experiment which underwent the cDNA synthesis procedure without the addition of reverse transcriptase. Only experiments with little or no amplification in the ‘no RT’ sample were considered suitable for further analysis. Gene-of-interest primers can be found in Table S10. Expression qPCR Ct-values were calculated and plotted, and statistical analyses were performed in RStudio.

### Staining, embedding, and sectioning

Plants expressing GUS reporters were stained with GUS staining solution (2 mM X-GlcA, 0.1 % v/v Triton X, 10 mM EDTA, 2.5 mM potassium hexacyanoferrate II, 2.5 mM potassium hexacyanoferrate III, 10.9 g/L sodium phosphate dibasic anhydrous, pH 7.4) at 37 °C for 12-16 hours. The GUS staining solution was then removed, samples rinsed with Phosphate Buffer Solution, then solution replaced with 70 % (v/v) ethanol. The samples were incubated in a 55 °C oven for at least 30 min, then the ethanol was poured off and replaced. This was repeated until the ethanol solution came off clear.

For sectioning of samples in promoter-reporter lines, plants were embedded in 3 % (w/v) agarose gel and cut by vibratome into 250-300 μm sections. This is the case for both *pro*Mp*GRAS7::tdTomato-NLS* and *pro*Mp*GRAS7::GUS* reporter lines, although the *pro*Mp*GRAS7::tdTomato-NLS* lines were embedded and cut as fresh tissue.

For histological observation of mutant antheridia, plants were fixed in a solution containing 3.7 % formaldehyde, 1 % (v/v) glutaraldehyde, 50 % (v/v) ethanol, 5 % (v/v) acetic acid, and 0.5 % (v/v)Tween 20, and left in a 4 °C fridge overnight. The samples were then dehydrated in an ethanol series, and infiltrated and embedded in LR white resin (Agar Scientific) according to the manufacturers’ instructions. Blocks were cut with a coping saw, and 1 μm sections were cut with glass knives. The samples were then stained with 0.1 % (w/v) Toluidine blue for observation of cellular structures.

### mRNA *in situ* Hybridization

The expression pattern of Mp*GRAS7* was analysed by mRNA in situ hybridization. A fragment of Mp*GRAS7* coding sequence was amplified using primers Mp*GRAS7*-ISH-P-F (5’-CGGTGAATTCCATCCTGCAG-3’) and Mp*GRAS7*-ISH-P-R (5’-

GCTACCGACGATGCATGTTT-3’). The PCR product was ligated into the pGEM-T Easy vector (Promega), and the orientation of the insertion was verified by sequencing. The construct was used as a template for PCR using T7 and SP6 primers. The PCR product was purified and used for *in vitro* transcription with the DIG RNA Labeling Kit (Roche).

Mature antheridiophores of *Marchantia polymorpha* were collected and fixed in FAA (3.7% v/v formaldehyde, 5% v/v acetic acid, 50% v/v ethanol). Samples were embedded in wax and cut into 8-µm sections. Slices were treated with Histoclear II (National Diagnostics, HS-202) for 10 min and in EtOH for 2 min, respectively. Histoclear II and EtOH treatments were repeated twice. Tissue sections were then rehydrated in 95 % EtOH, 1 min; 90 % EtOH, 1 min; 80 % EtOH, 1 min; 60 % EtOH/0.75 % NaCl, 1 min; 30 % EtOH/0.75 % NaCl, 1 min; 0.75 % NaCl, 2 min; and PBS, 2 min. After Proteinase K (19.2 mg/ml) digestion at 37 °C for 30 min, the reactions were stopped by glycine (2 mg/ml in PBS, 2 min), and then treated with FAA for 5 min, and PBS for 5 min. For dehydration, the samples were treated in 0.75 % NaCl, 2 min; 30 % EtOH/0.75 % NaCl, 30 s; 60% EtOH/0.75 % NaCl, 30 s; 80 % EtOH, 30 s; 90% EtOH, 30 s; 95% EtOH, 30 s; 100 % EtOH, 1 min; and 100 % EtOH fresh, 1 min.

Sections were hybridised with Mp*GRAS7* specific probes overnight at 55 °C. The probes bound to Mp*GRAS7* mRNAs were recognized by an anti-digoxigenin antibody (Roche). The hybridization signals were detected by the colour reaction using NBT/BCIP (Roche).

### Gemma cup emergence and dormancy assay

Plants were grown from gemmae, and gemma cups per plant were counted from 3-5 weeks post-plating. Gemma cups were viewed under a dissecting scope (Olympus stereoscope) and were determined to be dormant if there were no rhizoids growing from gemmae in the cup. Gemma cup dormancy index (GCDI) was calculated per plant by the number of gemma cups with rhizoids divided by the total number of gemma cups, so that 0=dormant and 1=not dormant.

### Gametangiophore emergence, FR, humidity shock, and ABA treatment experiments

In all FR light treatment experiments, plants were initially grown for two weeks in normal white light conditions (see plant growth conditions section), and then moved to conditions of white light supplemented with FR (wavelength 730-740 nm). Gametangiophore emergence was measured by counting the number of gametangiophores per plant at 1-week intervals after irradiation (1, 2, 3, and 4wpi). The time point used in the FR transcriptome experiment was 24 h post-irradiation. Humidity shock treatment consisted of placing open tissue culture plates in a laminar flow hood for 1 h, closing plates, and GUS staining after 4 h incubation. The ABA transcriptome experiment was carried out by droplet-treatment using 100 µM ABA, collecting at 1 and 4 hpt. Other ABA treatments were applied either through media or through droplet-treatment at the concentrations specified in figure legends.

### Transcriptomic analysis

RNA samples were derived, prepared, and sent to a commercial provider (Novogene UK) for mRNA library preparation and Illumina paired-end (PE150) sequencing at a sequencing depth of 30M paired reads per sample. Reads received were quality checked with FastQC (Babraham Bioinformatics) before and after trimming with Trimmomatic (Bolger *et al*., 2014) to remove adapters. There were no overrepresented sequences. Sample reads were then mapped to v6.1 of the *M. polymorpha* genome (Montgomery *et al*., 2020; available on *marchantia.info*, last accessed August 2023) using STAR (Dobin *et al*., 2014). Raw counts were obtained with featureCounts (Liao *et al*., 2014), and only uniquely mapped and properly paired reads were used for differential gene expression (DEG) analysis. DEG analysis was carried out using the DESeq2 (Bioconductor for *R*, Love *et al*., 2014) package. For each experiment, pairwise comparisons between mock vs treatment samples, and wild type vs mutant were carried out. Nomenclature tables for *M. polymorpha* genes were downloaded from *marchantia.info* and combined with DEG tables using the ‘full_join’ function (‘dplyr’ package) in *R* (version 4.1.0). Log_2_FC values and *p*-adj values used as cut-offs are included in figure legends.

### Phylogenetic tree construction and gene presence/absence assessment

Input sequences were aligned using MAFFT (Katoh *et al*., 2013). Sequences were trimmed using TrimAL on the ‘gappyout’ setting (Capella-Gutiérrez *et al*., 2009). The aligned and trimmed sequences were then run through IQ-TREE 2 (Minh et al., 2020). Output tree files were visualised using iTOL (Letunic & Bork, 2021).

SCLA sequences were determined by construction of phylogenetic trees of all GRAS families such that the functional sub-clades resolve. Sequences from Cenci & Rouard (2017) were used to assign ortho-groups. Gene presence in the SCLA group was then determined by looking at individual ortho-groups, and absence of gene families was inferred from this assessment. The sequences used for phylogeny in Fig. 1 are listed in Table S1A.

GRAS protein sequences were collected from publicly available databases (Plant TFDB, Jin *et al*., 2016; iTAK, Zheng *et al*., 2016). For genomes which were not available on Plant TFDB or iTAK, sequences for relevant proteins were determined by HMMER3 software using the phmmer function (Eddy, 2011), using a protein from a related species as an input and a proteome FASTA file as the database.

### Promoter analysis

The upstream promoter regions of SCLA orthologs used were limited to genomes available on both Phytozome (Goodstein *et al*., 2011) and PlantTFDB (Jin *et al*., 2016), with the exceptions of *Marchantia paleacea*, *Anthoceros agrestis*, *Ginkgo biloba* and *Nicotiana benthamiana*, which were manually included due to their experimental or evolutionary relevance. Sequences were extracted from Phytozome where possible and sequences extracted directly from genomes manually when necessary. 3 kb sequences upstream of the start site were taken to represent the promoters (sequences available Table S1B). Putative transcription factor binding sites were predicted using the PlantRegMap binding site prediction online tool (Tian *et al*., 2019). Only binding sites with *p*-values of <0.001 were used. Promoter sequences were manually annotated for Fig. S4A-C.

### Statistical analysis and quantification

Details of statistical tests and n-numbers used for each experiment can be found in figure legends. Tukey’s Honest Statistical Difference (HSD) post-hoc analysis for statistical groupings was carried out in *R* (version 4.1.0; dependency: ‘agricolae’; Mendiburu and Yaseen, 2020). Exact *p*-value tests can also be found in figure legends. For transcriptomic data, the cutoff used was *p-adj*<0.01 unless otherwise stated. t-tests (for non-transcriptomic data) were carried out in *Microsoft excel*.

## Supplementary Information

Supplementary Tables S1A-P

Table S1A GRAS Sequences determined to be SCLA-type by phylogenetics

Table S1B SCLA promoter sequences used for cross-taxa promoter analysis

Table S1C DEGs WL vs FR (Tak-1) Treatment comparisons Log2FC=>2 or =<-2; p-adj=0.01

Table S1D DEGs WL vs FR (*gras7-1*) Treatment comparisons Log2FC=>2 or =<-2; p-adj=0.01

Table S1E DEGs mock vs ABA (Tak-1 1 hpt) Treatment comparisons Log2FC=>2 or =<-2; p-adj=0.01

Table S1F DEGs mock vs ABA (*gras7-1* 1 hpt) Treatment comparisons Log2FC=>2 or =<-2; p-adj=0.01

Table S1G DEGs mock vs ABA (Tak-1 4 hpt) Treatment comparisons Log2FC=>2 or =<-2; p-adj=0.01

Table S1H DEGs mock vs ABA (*gras7-1* 4 hpt) Treatment comparisons Log2FC=>2 or =<-2; p-adj=0.01

Table S1I DEGs Tak-1 vs *gras7* (WL) Genotype comparisons (Log2FC=>1 or =<-1; p-adj=0.01)

Table S1J DEGs Tak-1 vs *gras7* (FR) Genotype comparisons (Log2FC=>1 or =<-1; p-adj=0.01)

Table S1K DEGs which are FR-responsive (p-adj=0.01) in WT and significantly differentially expressed between WT and gras7 (Log2FC=>1 or =<-1, p-adj=0.01).

Table S1L DEGs Tak-1 vs *gras7* (mock 1 hpt) Genotype comparisons (Log2FC=>1 or =<-1; p-adj=0.01)

Table S1M DEGs Tak-1 vs *gras7* (mock 4 hpt) Genotype comparisons (Log2FC=>1 or =<-1; p-adj=0.01)

Table S1N DEGs Tak-1 vs *gras7* (ABA 1 hpt) Genotype comparisons (Log2FC=>1 or =<-1; p-adj=0.01)

Table S1O DEGs Tak-1 vs *gras7* (ABA 4 hpt) Genotype comparisons (Log2FC=>1 or =<-1; p-adj=0.01)

Table S1P DEGs which are ABA-responsive (p-adj=0.01) in WT and significantly differentially expressed between WT and gras7 at either 1 hpt or 4 hpt (Log2FC=>1 or =<-1, p-adj=0.01).

Table S2 Constructs used in this study

Table S3 Constructs produced in this study

Table S4 Cloning primers

Table S5 sgRNA sequences

Table S6 Oligos compatible with pMpGE_En01 vector

Table S7 Experimental Models: Organisms/Strains

Table S8 Genotyping primers

Table S9 PCR cycle details

Table S10 qPCR primers

