## Supplementary Figures and Tables S2-10 for "An environment-responsive SCLA-type GRAS protein controls reproductive strategy in *Marchantia polymorpha*"

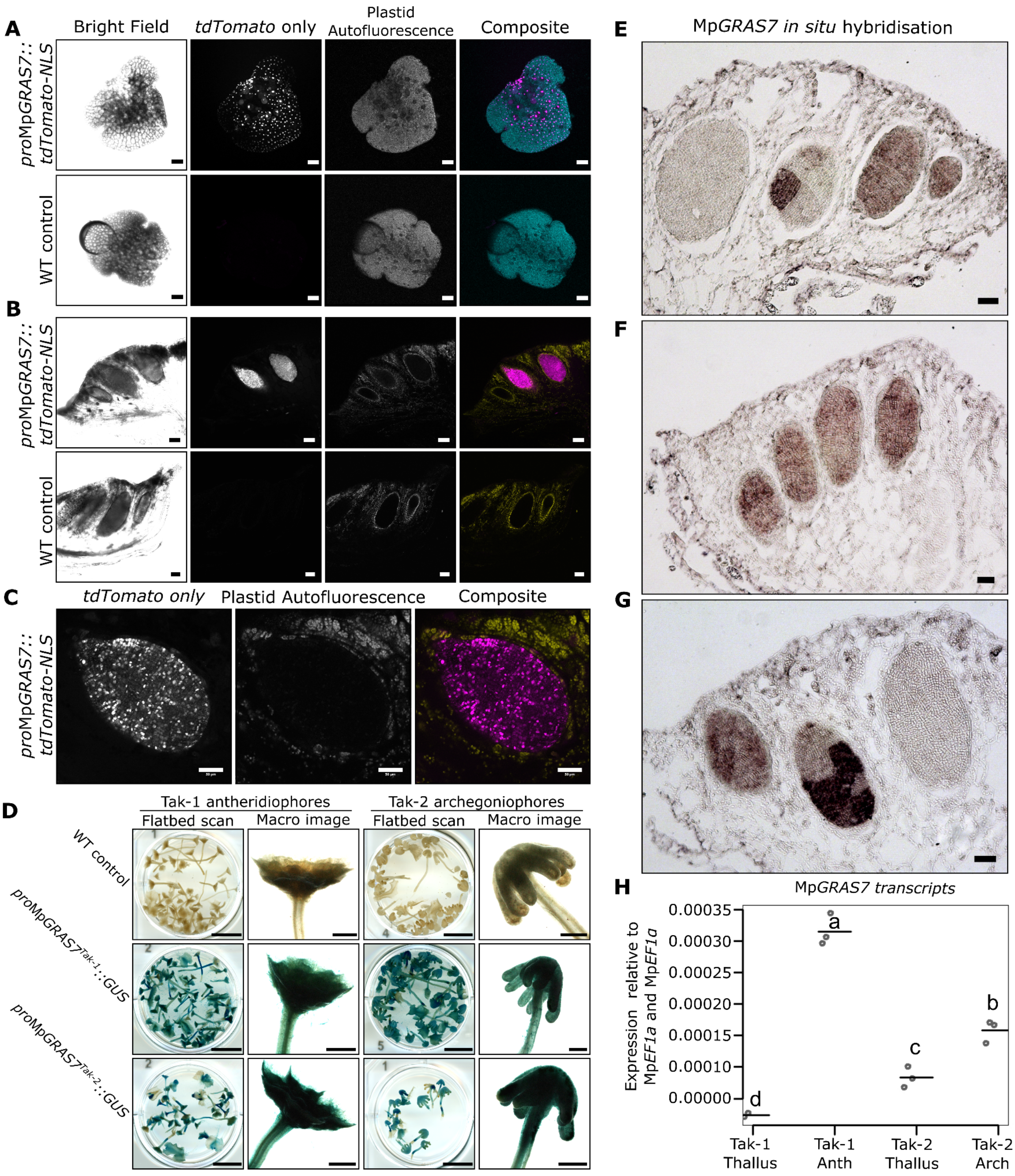


**Supplementary Figure 1 (relating to Fig. 1). Additional data on Mp*GRAS7* tissue/organ-specific expression.** Confocal imaging of the *pro*Mp*GRAS7::tdTomato-NLS* reporter in gemmae **(A)** and antheridia **(B)**, scale bars = 100 µm. tdTomato signal shown in magenta, plastid autofluorescence shown in cyan in A, yellow in B. **C)** 25x objective confocal image of an antheridium expressing *pro*Mp*GRAS7:tdTomato-NLS* with clear expression in developing sperm cells. Scale bars = 50 µm. **D)** Flatbed scan and light microscopy (macro image) images of late-stage (10 weeks post-induction) gametangiophores expressing GUS under the *pro*Mp*GRAS7* cloned from either Tak-1 (*pro*Mp*GRAS7*$^{Tak\text{-}1}$) or Tak-2 (*pro*Mp*GRAS7*$^{Tak\text{-}2}$) gDNA transformed into either *Tak-1* or *Tak-2* WT backgrounds. Previously tested *pro*Mp*GRAS7*$^{Tak\text{-}1}$*::GUS* plants were used as a positive control and WT plants were used as a negative control. 1x line of *pro*Mp*GRAS7*$^{Tak\text{-}1}$*::GUS/*Tak-1 plants were tested for activation in the antheridiophores (41/41 expressed GUS in the antheridiophore), 1x line of *pro*Mp*GRAS7*$^{Tak\text{-}1}$*::GUS/Tak-2* plants were tested for activation in the archegoniophores (31/31), 4x independent transgenic lines of *pro*Mp*GRAS7*$^{Tak\text{-}2}$*::GUS/*Tak-1 plants were tested for activation in the antheridiophores (70/74), 4x independent transgenic lines of *pro*Mp*GRAS7*$^{Tak\text{-}2}$*::GUS/Tak-2* plants were tested for activation in the archegoniophores (44/44). Images shown are of single transgenic lines only. Scale bars in flatbed scan images = 1cm, scale bars in macro images = 1000 µm. **E-G)** additional images of Mp*GRAS7 in situ* hybridisation in antheridia. Scale bars = 100 µm. **H)** RT-qPCR data of Mp*GRAS7* transcripts comparing thallus tissue against gametangiophore tissue in WT Tak-1 and Tak-2 lines. *n=3*, each data point is a biological replicate, with each sample containing 3-4 individuals harvested per biological sample. qPCR analysis was carried out in triplicate. Statistical groupings are Tukey's HSD Post-hoc analysis ($p$<0.05).

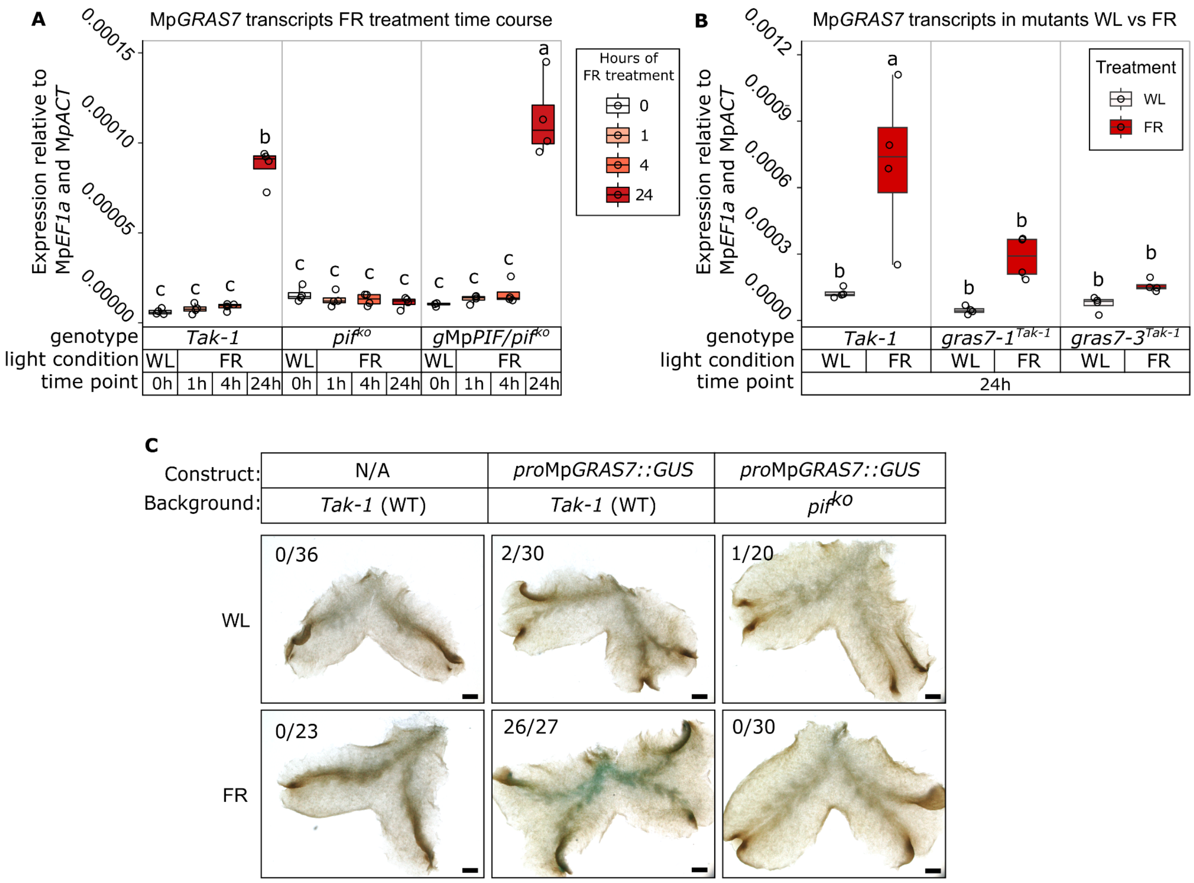


**Supplementary Figure 2. Mp*GRAS7* is responsive to FR treatment in a Mp*PIF*-dependent manner. A)** RT-qPCR experiment of Mp*GRAS7* transcripts during a FR treatment time course. Samples are grouped by genotype and time point. The 0h treatment samples were not treated with FR light (white light, WL, only). **B)** Additional RT-qPCR experiment of MpGRAS7 transcripts after 24 hours FR treatment. WL samples were harvested at the same time. For A-B, *n*=4, each data point is a biological replicate, with each sample containing 3-4 individuals harvested per biological sample. RT-qPCR reactions carried out in triplicate. Statistical groupings shown are from Tukey's HSD post-hoc analysis. **C)** Light microscopy images of GUS stained plants at the 24h FR treatment time point. This full GUS stain experiment is shown partially in Fig. 2A. WL = white light control, plants untreated with FR light. *pro*Mp*GRAS7::GUS* constructs were transformed into WT *Tak-1* and *pif*$^{ko}$ backgrounds and two independent transgenic lines were tested. Scale = 1000 µm.

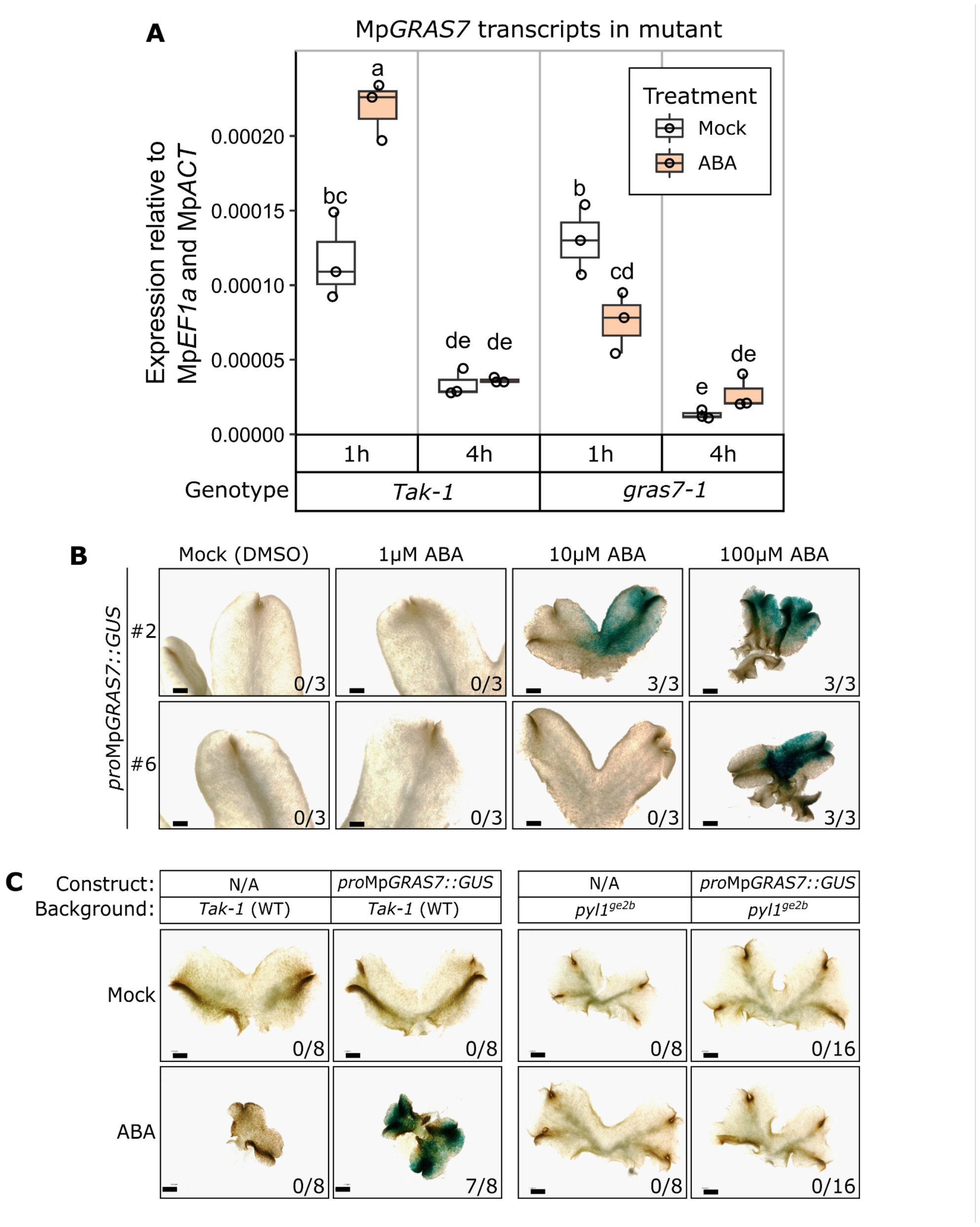


**Supplementary Figure 3. Mp*GRAS7* is responsive to ABA treatment in a Mp*PYL1*-dependent manner. A)** RT-qPCR experiment of Mp*GRAS7* transcripts during dropwise ABA (100 μM) treatment including *gras7-1* mutant lines. *n*=3, each data point is a biological replicate, with each sample containing 3-4 individuals harvested per biological sample. This experiment was used for transcriptome sequencing. RT-qPCR analysis was carried out in triplicate. Statistical tests shown for A-B are Tukey's HSD post-hoc analysis ($p$<0.05). **B)** Two additional transgenic lines of *pro*Mp*GRAS7::GUS* subjected to mock (low concentration of DMSO, equivalent to the 100 μM ABA treatment) treatment or ABA treatment via growth media, supplementary to Fig. 2B. Scale = 1000 μm. **C)** Light microscopy images of a GUS experiment of 2-week old plants grown on mock media (top row) or 100 μM ABA (bottom row). *n*=8 individuals per genotype per treatment, numbers in the corners of the pictures indicate the number of individuals which showed any blue GUS staining. Super-transformed mutant *pyl1*$^{ge2b}$ was tested using two independent transgenic lines and stained alongside a mutant not containing the GUS construct. Scale = 1000 μm.

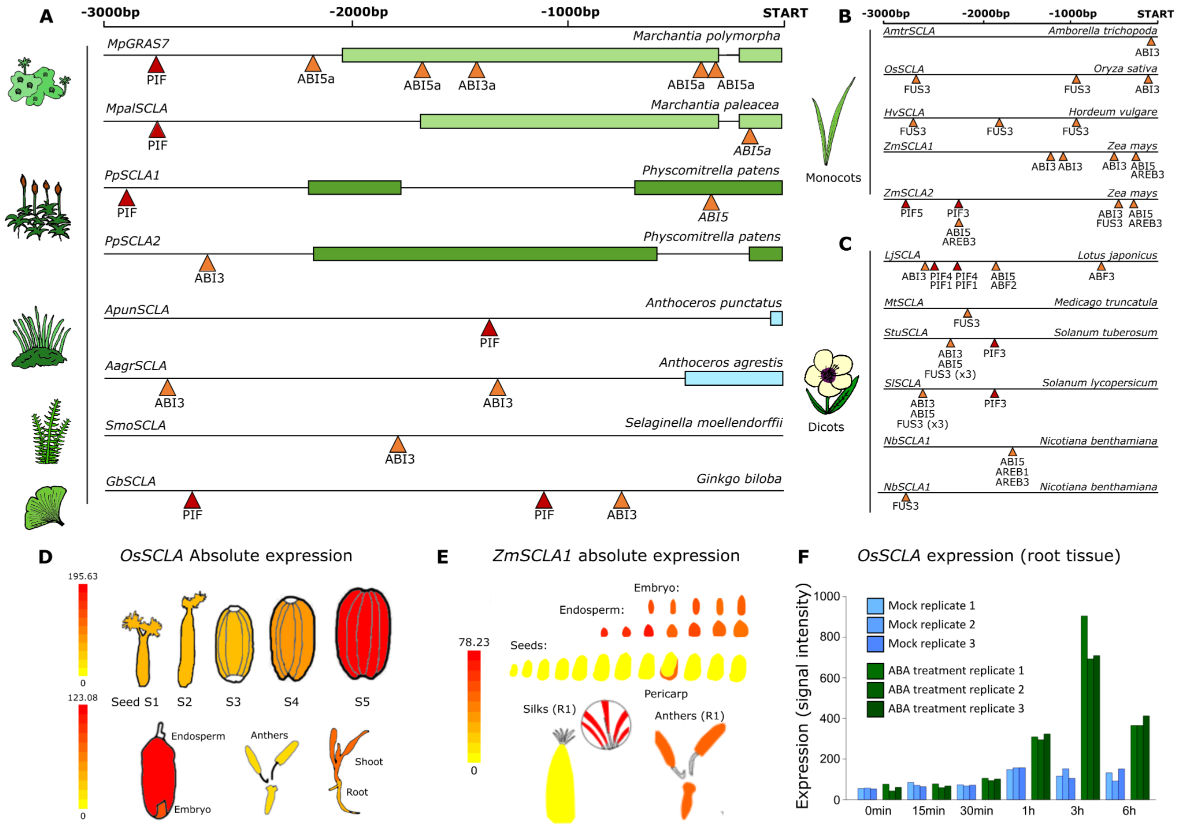

**Supplementary Figure 4. Cis-elements and public transcriptomic data from SCLA sub-family members in land plants.**

**A)** SCLA promoters in non-seed plants: *Marchantia polymorpha* Mp*GRAS7* (Mp8g01770.1), *Marchantia paleacea* Mpa*SCLA* (Marpal_utg000030g0064951), *Physcomitrium patens* Pp*SCLA1,* Pp*SCLA2* (Pp1s165_77V6), *Anthoceros punctatus* Apun*SCLA* (Apun_evm.model.utg000126l.90.1), *Anthoceros agrestis* Aagr*SCLA* (AagrOXF_evm.model.utg000009l.366.1), *Selaginella moellendorffii* Smo*SCLA* (102631), *Ginkgo biloba* Gb*SCLA* (Gb_20415) **B)** SCLA promoters in monocots: *Amborella trichopoda* Amtr*SCLA* (AMBTR_LOC18436680;AmTrH1.11G080100.1; evm_27.model.AmTr_v1.0_scaffold00017.27), Oryza sativa Os*SCLA* (LOC_Os04g35250), *Hordeum vulgare* Hv*SCLA* (HORVU.MOREX.r3.2HG0168770.1), *Zea mays* Zm*SCLA1* (GRMZM2G176537_P01), and Zm*SCLA2* (GRMZM2G420280_P01). **C)** SCLA promoters in dicots: *Lotus japonicus* Lj*SCLA* (Lj5g0014151.1), *Medicago truncatula* Mt*SCLA* (Medtr1g096030.1), *Solanum tuberosum* Stu*SCLA* (PGSC0003DMP400016289), *Solanum lycopersicum* Sl*SCLA* (Solyc11g017100.1.1), *Nicotiana benthamiana* Nb*SCLA1* (Niben101Scf01365g06013.1), and Nb*SCLA2* (Niben101Scf03473g01018.1). For A-C, putative binding sites were predicted using PlantRegMap. ABA responsive elements are shown in orange, PIF binding sites are shown in red. Boxes in promoters indicate 5’ UTR regions. Sequences used for analysis can be found in Table S1B. **D)** Tissue-level expression patterns of rice ortholog Os*SCLA* in publicly available expression data, expression shown is absolute. **E)** Tissue level expression patterns of *Zea mays* ortholog Zm*SCLA1* in publicly available expression data, expression shown is absolute. Zm*SCLA2* expression patterns are similar. For D-E, transcriptomic data was taken from ePlant. (Waese *et al*. 2017). **F)** Os*SCLA* root tissue expression in response to treatment with abscisic acid, data taken from RiceXPro transcriptome (Sato *et al.* 2013). Colours of the original graph have been changed to be colour-blind friendly.

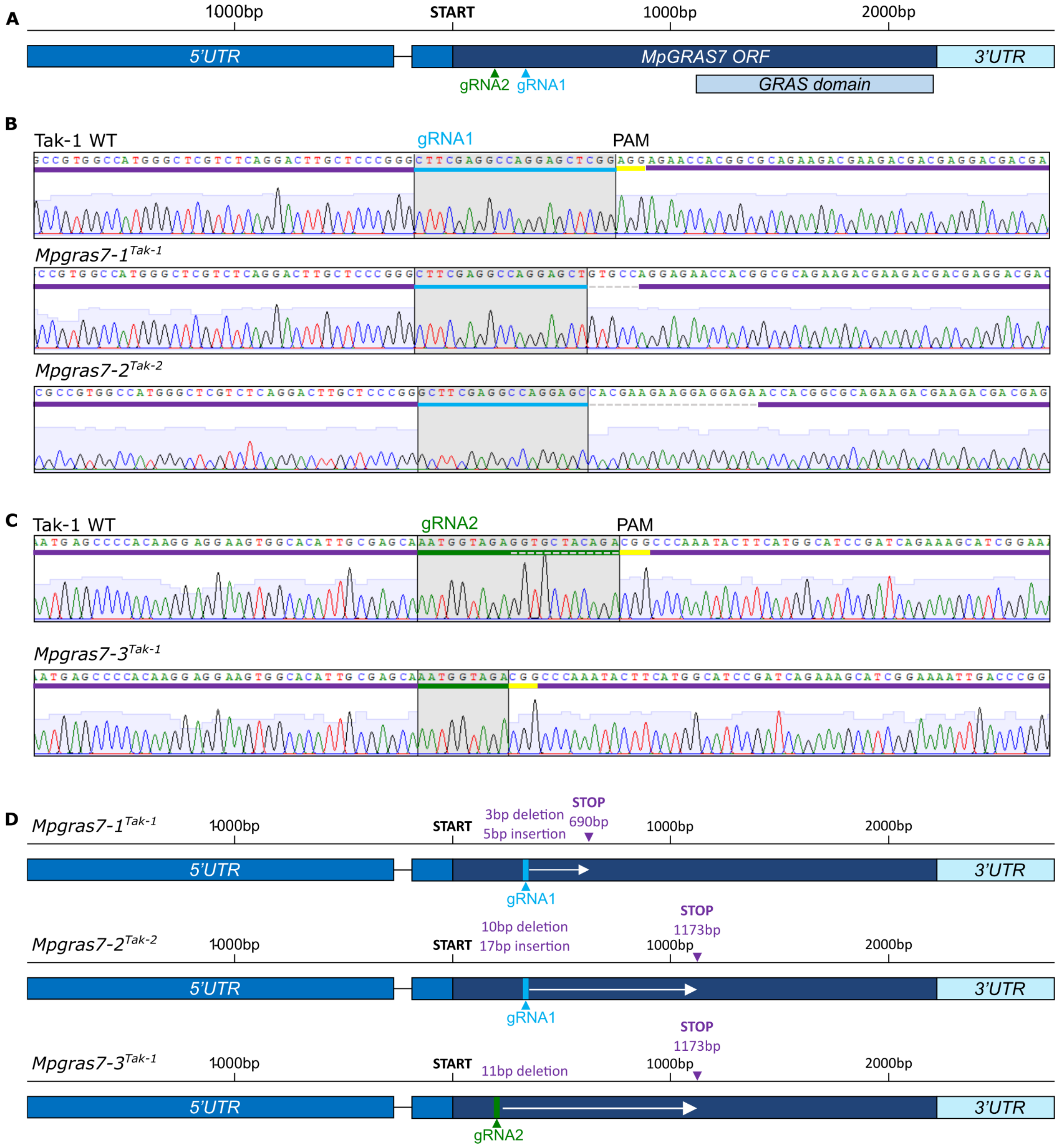


**Supplementary Figure 5. CRISPR/Cas9-induced mutations in Mp*GRAS7*. A)** Graphical representation of the open reading frame of Mp*GRAS7*, gRNA1 is indicated in cyan, gRNA2 indicated in green. The 5' UTR is indicated blue, the open reading frame (ORF) is indicated in dark blue, and the 3' UTR is indicated in light blue. The grey-blue box shows the GRAS domain. **B)** Sanger sequencing results of the region targeted by gRNA1 (blue). Alleles in both male (Mp*gras7-1*$^{Tak\text{-}1}$) and female (Mp*gras7-2*$^{Tak\text{-}2}$) backgrounds were maintained for this sgRNA. gRNA-adjacent PAM sites are shown in yellow, and the indel mutations are indicated with a grey dashed line. **C)** Sanger sequencing results of the region targeted by sgRNA2 (green). The region deleted in Mp*gras7-3*$^{Tak\text{-}1}$ is indicated on the WT sequence by a grey dashed line. **D)** Graphical representations of the three mutant alleles used in this study, and the putative impacts on protein synthesis. All three mutations introduce frameshift mutations and early stop codons.

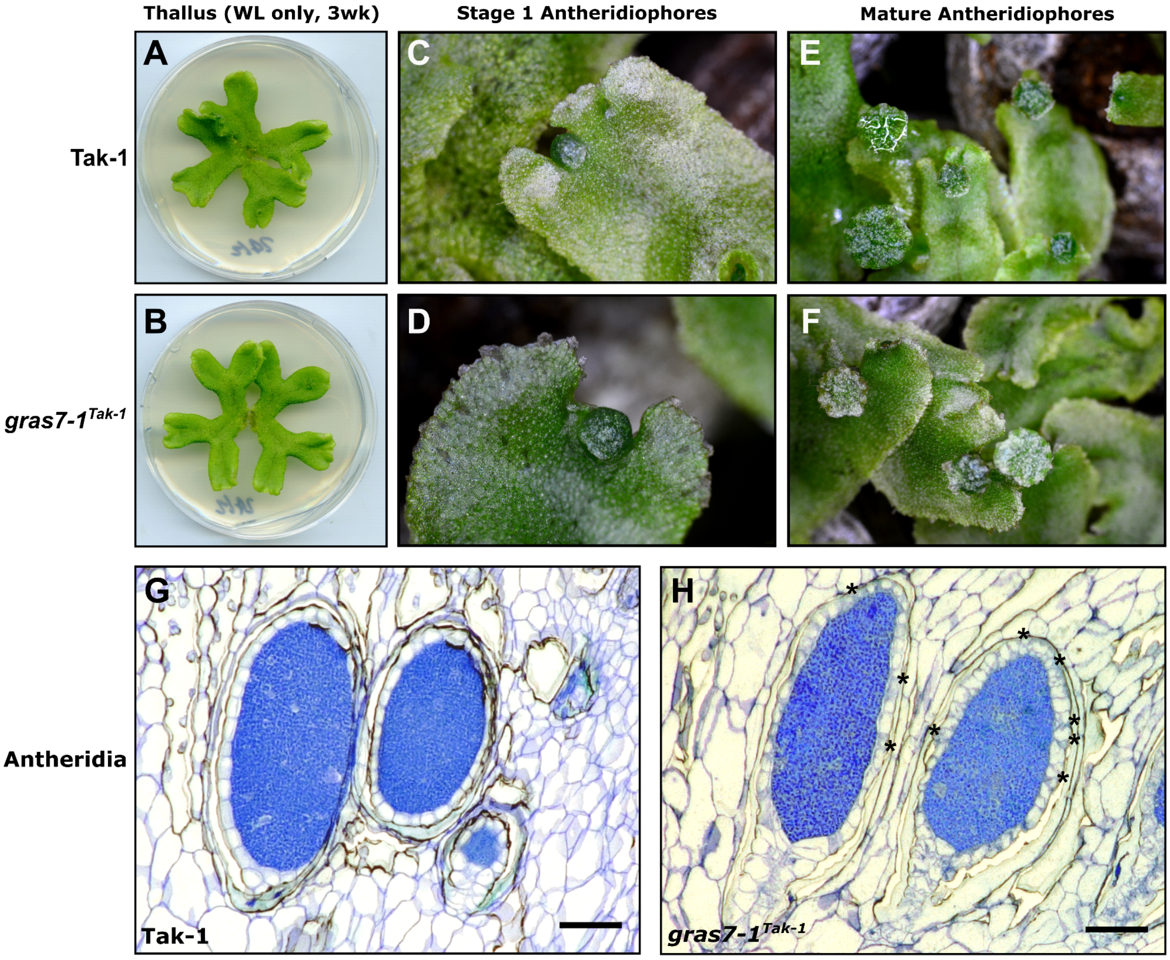


**Supplementary Figure 6. Growth of WT and *gras7-1$^{Tak-1}$* mutant male reproductive tissue. A)** Tak-1 and **B)** *gras7-1$^{Tak-1}$* thallus tissue grown from gemmae on ½ MS B5 (pH 6.7), images taken 3 weeks post plating. **C)** Tak-1 and **D)** *gras7-1$^{Tak-1}$* stage 1 antheridiophores. **E)** Tak-1 and **F)** *gras7-1$^{Tak-1}$* mature antheridiophores. **G)** Tak-1 and **H)** *gras7-1$^{Tak-1}$* antheridia. 5μm sections from samples embedded in Historesin, stained with 0.1% Toluidine Blue, imaged on a Keyence VHX-5000 digital microscope. Asterisks indicate jacket cells with aberrant cell patterning. Scale bar = 100μm.

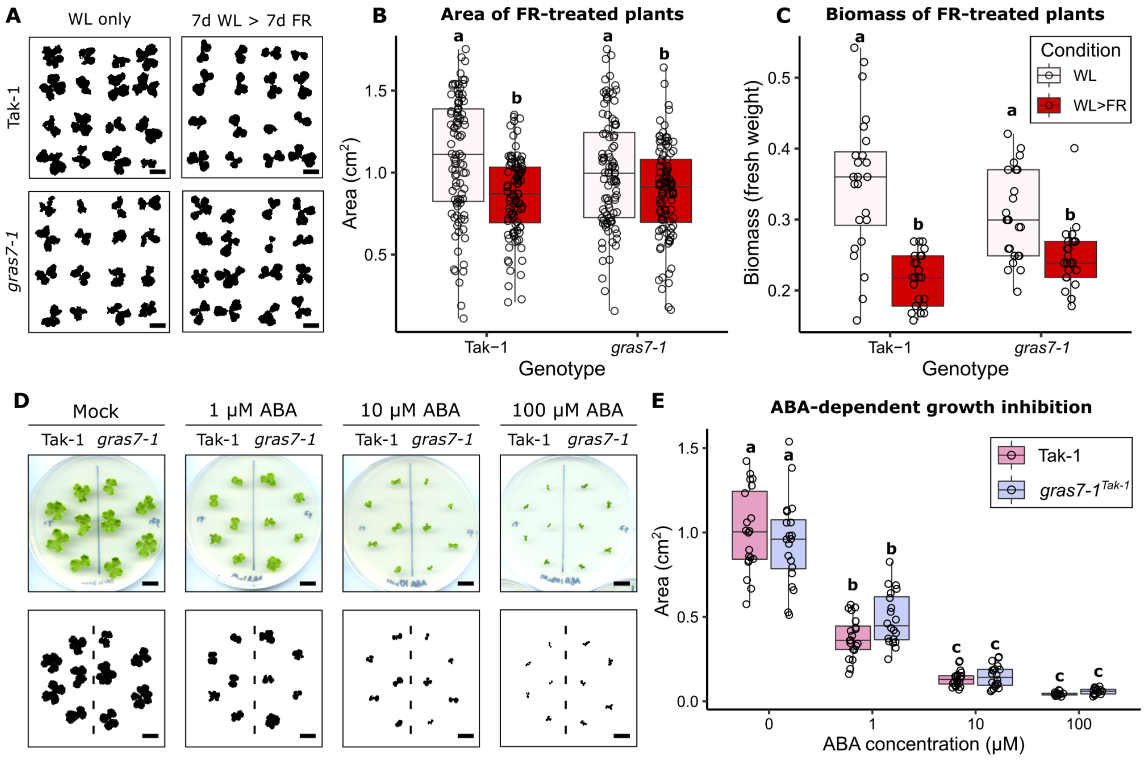


**Supplementary Figure 7. Mp*gras7*$^{ge}$ plants are unaltered in growth inhibition response in ABA treatments when compared to WT Tak-1. A)** Threshold images of the blue channel of plants used for measuring area ($cm^2$). Plants are treated with either white light (WL) only, or WL for 7 days, followed by 7 days of high FR irradiation. Scale bar = 1 cm. **B)** Area of WT Tak-1 and *gras7-1* plants when exposed to WL only or WL then FR. $n$=16 per replicate, $N$=6 replicates. **C)** Biomass (fresh weight in grams) of WT Tak-1 and *gras7-1* plants when exposed to WL only or WL then FR. Each data point is the combined weight of 4 individuals. $n$=4 per replicate, $N$=6 replicates. **D)** Plants grown on increasing concentrations of ABA. First row are raw flatbed scan images, second row are sample threshold images from the blue channel, used for measuring area. Scale bar = 1 cm. **E)** Quantification of ABA-dependent growth inhibition comparing WT and *gras7-1*$^{Tak\text{-}1}$ area. $n$=5 plants per condition per genotype, $N$=3 replicates. Areas measured on ImageJ, statistical analyses performed are Tukey's HSD Post-hoc analysis ($p<0.05$).

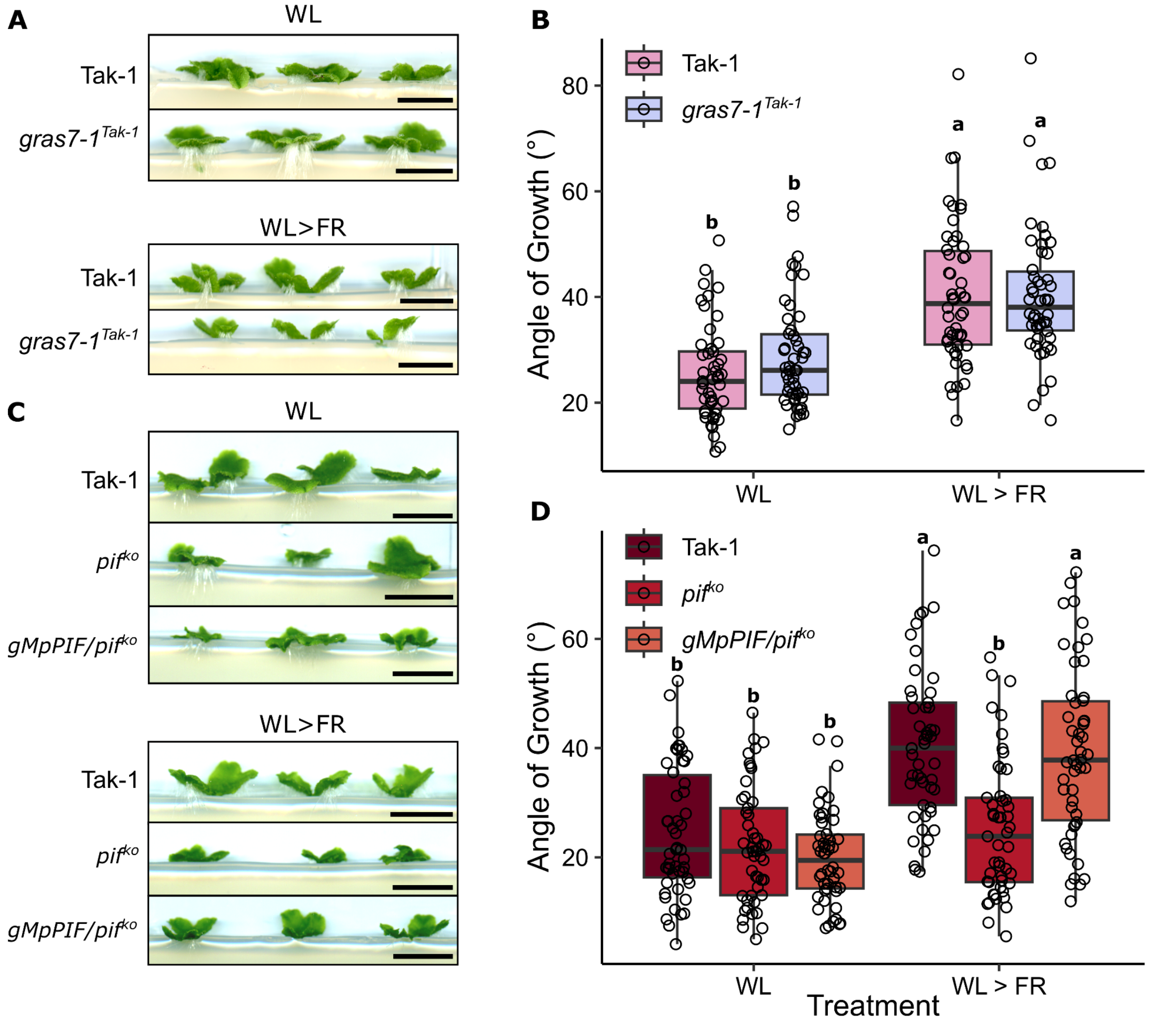


**Supplementary Figure 8. Mp*gras7*$^{ge}$ mutants are not affected in orthotropic growth FR-response, whereas Mp*pif*$^{ko}$ mutants are. A)** Orthotropic growth of WT Tak-1 and *gras7-1*$^{Tak\text{-}1}$ in constant White Light (WL) or White Light moved to White Light supplemented with high FR light (WL>FR) conditions. *gras7-1*$^{Tak\text{-}1}$ mutants are WT-like in this condition. Scale bars = 10mm. **B)** Quantification of the angle of growth of Tak-1 WT vs *gras7* mutants in WL or WL>FR conditions. **C)** Orthotropic growth of Mp*pif*$^{ko}$ mutants, the corresponding Tak-1 WT*,* and the complemented line *g*Mp*PIF/*Mp*pif*$^{ko}$ in WL or WL>FR conditions. Scale bars = 10mm. **D)** Quantification of the angle of growth of Mp*pif*$^{ko}$ mutants, the corresponding Tak-1 WT*,* and the complemented line *g*Mp*PIF/*Mp*pif*$^{ko}$ in WL or WL>FR conditions. Angles measured in ImageJ, statistical groupings are Tukey's HSD Post-hoc analysis ($p$<0.05). $n$=5 plants per replicate, $N$=5 replicates. Each individual plant yields 2 angle measurements, total ~50 data points per genotype per condition.

### Supplementary Tables S2-S10

#### Supplementary Table 2: Constructs used in this study

| Construct | Addgene entry | Source |
|---|---|---|
| pDONR-221 | https://www.addgene.org/vector-database/2394/ | Our laboratory collection |
| pMpGWB104 | https://www.addgene.org/68558/ | Ishizaki *et al*. (2015) |
| pMpGWB316 | https://www.addgene.org/68644/ | Ishizaki *et al*. (2015) |
| pMpGWB304 | https://www.addgene.org/68632/ | Ishizaki *et al*. (2015) |
| pMpGWB105 | https://www.addgene.org/68559/ | Ishizaki *et al*. (2015) |
| pMpGWB106 | https://www.addgene.org/68560/ | Ishizaki *et al*. (2015) |
| pMpGE_En01 | https://www.addgene.org/71534/ | Sugano *et al*. (2018) |
| pMpGE010 | https://www.addgene.org/71536/ | Sugano *et al*. (2018) |
| pMpGWB301 | https://www.addgene.org/68629/ | Sugano *et al*. (2018) |

#### Supplementary Table 3: Constructs produced in this study

| Construct name | Description |
|---|---|
| *proMpGRAS7-221* | Mp*GRAS7* promoter in pDONR-221 backbone. |
| *MpGRAS7-5'UTR-221* | Mp*GRAS7* 5' UTR only in pDONR-221 backbone. |
| *proMpGRAS7-pMpGWB104* | Mp*GRAS7* promoter in a destination vector driving *GUS*. Hygromycin resistance. |
| *proMpGRAS7-pMpGWB316* | Mp*GRAS7* promoter in a destination vector driving *tdTomato-NLS*. Chlorsulfuron resistance. |

| | |
|---|---|
| *proMpGRAS7-pMpGWB304* | Mp*GRAS7* promoter in a destination vector driving *GUS*. Chlorsulfuron resistance. |
| *MpGRAS7-5'UTR-pMpGWB104* | Mp*GRAS7* 5' UTR only in a destination vector driving *GUS*. Hygromycin resistance. |
| *MpGRAS7 sgRNA1 (SacI)-En01* | Mp*GRAS7* sgRNA1 cloned into pMpGE_En01 gene editing entry vector. |
| *MpGRAS7 sgRNA1 (SacI)-GE010* | Mp*GRAS7* sgRNA1 cloned into pMpGE010 destination vector. Hygromycin resistance. |
| *MpGRAS7 sgRNA2 (SfcI)-En01* | Mp*GRAS7* sgRNA2 cloned into pMpGE_En01 gene editing entry vector. |
| *MpGRAS7 sgRNA2 (SfcI)-GE010* | Mp*GRAS7* sgRNA2 cloned into pMpGE010 destination vector. Hygromycin resistance. |
| *ccdb-synGRAS7-pUC* | ccdb/CmR cassette upstream of synthetic GRAS7 in a pUC backbone. |
| *ccdb-synGRAS7-301* | ccdb/CmR cassette upstream of synthetic GRAS7 in a Marchantia expression vector. Chlorsulfuron resistance. |
| *proMpGRAS7::GRAS7* | Native promoter driving synthetic GRAS7 in a Marchantia expression vector. Chlorsulfuron resistance. |

**Supplementary Table 4: Cloning primers**

| **Primer Name** | **Sequence** | **Source** |
|---|---|---|
| *MpGRAS7pro-B1* | AAAAAGCAGGCTATGAAGCTGGAATAAATCGGCTG | This study |
| *MpGRAS7pro-B2* | AGAAAGCTGGGTCGGCGGCTACCCTGCTCGCTCTC CCG | This study |
| *MpGRAS7-UTR-B1* | AAAAAGCAGGCTGGACGAAGGGCGAGTGTGAGG | This study |

**Supplementary Table 5: sgRNA sequences**

| **sgRNA Name** | **Sequence** | **Source** |
|---|---|---|
| *MpGRAS7-gRNA1-SacI* | CTTCGAGGCCAGGAGCTCGG | This study |

| | | |
|---|---|---|
| *MpGRAS7-gRNA2-SfcI* | AATGGTAGAGGTGCTACAGA | This study |

**Supplementary Table 6: Oligos compatible with pMpGE_En01 vector**

| **Primer Name** | **Sequence** | **Source** |
|---|---|---|
| *MpGRAS7-gRNA1-SacI-F* | GCACCCAGCCTCTCGCTTCGAGGCCAGGA GCTCGGGTTTTAGAGCTAGAA | This study |
| *MpGRAS7-gRNA1-SacI-R* | TTCTAGCTCTAAAACCCGAGCTCCTGGCCT CGAAGCGAGAGGCTGGGTGC | This study |
| *MpGRAS7-gRNA2-SfcI-F* | GCACCCAGCCTCTCGAATGGTAGAGGTGCT ACAGAGTTTTAGAGCTAGAA | This study |
| *MpGRAS7-gRNA2-SfcI-R* | TTCTAGCTCTAAAACTCTGTAGCACCTCTAC CATTCGAGAGGCTGGGTGC | This study |

**Supplementary Table 7: Experimental Models: Organisms/Strains**

| **Organism/strain** | **Source** |
|---|---|
| *Marchantia polymorpha* Tak-1, Tak-2 | Prof. Jim Haseloff, University of Cambridge |
| *Marchantia polymorpha* Cam-1, Cam-2 | Prof. Jim Haseloff, University of Cambridge |
| *pro*Mp*GRAS7::GUS* | This study |
| Mp*GRAS7-5'UTR::GUS* | This study |
| Mp*gras7-1*$^{Tak\text{-}1}$ | This study |
| Mp*gras7-2*$^{Tak\text{-}2}$ | This study |
| Mp*gras7-3*$^{Tak\text{-}1}$ | This study |
| *proMpGRAS7::GRAS7/ gras7-1*$^{Tak\text{-}1}$ | This study |
| *Marchantia polymorpha* Tak-1 (Kohchi) | Jahan *et al*. (2019) |

| | |
|---|---|
| *pyl1*$^{ge2b}$ | Jahan *et al*., (2019) |
| *Marchantia polymorpha* Tak-1 (HG) | Hernandez-Garcia *et al*. (2021) |
| *pif*$^{ko}$ | Hernandez-Garcia *et al*. (2021) |
| *gMpPIF/pif*$^{ko}$ | Hernandez-Garcia *et al*. (2021) |
| *pro*Mp*GRAS7::GUS/pyl1*$^{ge2b}$ | This study |
| *pro*Mp*GRAS7::GUS/pif*$^{ko}$ | This study |

**Supplementary Table 8: Genotyping primers**

| Primer Name | Sequence | Source |
|---|---|---|
| Mp*GRAS7*-F | GCGTAGTGTTGGAGCGAGAATTT | This study |
| Mp*GRAS7*-R | GTCGTCCGACTCCAGCATGCTG | This study |
| MpGRAS7-R2 | CTGAGCAACGAGCCCAACTGC | This study |
| *att*B1 universal primer | GGGGACAAGTTTGTACAAAAAAGCAGGCT | Gateway® user manual |
| *att*B2 universal primer | GGGGACCACTTTGTACAAGAAAGCTGGGT | Gateway® user manual |

**Supplementary Table 9: PCR cycle details**

| Primer Pair | Polymerase enzyme(s) used | Initial [Denaturing Temp] | Cycles (x39) | | | Final 72 °C |
|---|---|---|---|---|---|---|
| | | | [Denaturing Temp] °C | [Annealing Temp] °C | 72 °C | |
| Mp*GRAS7*-F, Mp*GRAS7*-R | Phusion,<br>Phire direct | 98 °C, 30 s | 98 °C, 20 s | 60 °C, 15 s | 7 s | 5 min |
| Mp*GRAS7*-F, Mp*GRAS7*-R | Gotaq | 95 °C, 2 min | 95 °C, 30 s | 60 °C, 30 s | 30 s | 5 min |

| | | | | | | |
|---|---|---|---|---|---|---|
| *att*B1uni, *att*B2uni | Phire direct | 98 °C, 5 s | 98 °C, 5 s | 60 °C, 5s | 98 °C, 20 s per 1 kb | 5 min |

**Supplementary Table 10: qPCR primers**

| Primer Name | Sequence | Source |
|---|---|---|
| Mp*ACT*-qF | AGGCATCTGGTATCCACGAG | Saint-Marcoux *et al.*, 2015 |
| Mp*ACT*-qR | AGGCATCTGGTATCCACGAG | Saint-Marcoux *et al.*, 2015 |
| Mp*EF1a*-qF | CCGAGATCCTGACCAAGG | Saint-Marcoux *et al.*, 2015 |
| Mp*EF1a*-qR | GAGGTGGGTACTCAGCGAAG | Saint-Marcoux *et al.*, 2015 |
| Mp*GRAS7*-qF | AGCGAGAATTTCCCAAGTACAC | This study |
| Mp*GRAS7*-qR | CCGTCTGTAGCACCTCTACCAT | This study |
| Mp*NCED*-qF | GGACTGCTTCTGCTTTCACC | This study |
| Mp*NCED*-qR | CGATCGCCAGGTACAAGAAT | This study |
